# Epac1 regulates cellular SUMOylation by promoting the formation of SUMO-activating nuclear condensates

**DOI:** 10.1101/2022.01.12.476066

**Authors:** Wenli Yang, William G. Robichaux, Fang C. Mei, Wei Lin, Li Li, Sheng Pan, Mark A. White, Yuan Chen, Xiaodong Cheng

**Affiliations:** Department of Integrative Biology and Pharmacology, The University of Texas Health Science Center, Houston, Texas, USA; Brown Foundation Institute of Molecular Medicine, The University of Texas Health Science Center, Houston, Texas, USA; Sealy Center for Structural Biology and Molecular Biophysics, The University of Texas Medical Branch, Galveston, Texas, USA; Department of Biochemistry and Molecular Biology, The University of Texas Medical Branch, Galveston, Texas, USA; Department of Surgery and Moores Cancer Center, UC San Diego Health, La Jolla, California, USA

## Abstract

Protein SUMOylation plays an essential role in maintaining cellular homeostasis when cells are under stress. However, precisely how SUMOylation is regulated, and a molecular mechanism linking cellular stress to SUMOylation remains elusive. Herein, we report that cAMP, a major stress-response second messenger, acts through Epac1 as a regulator of cellular SUMOylation. The Epac1-associated proteome is highly enriched with components of the SUMOylation pathway. Activation of Epac1 by intracellular cAMP triggers phase separation and the formation of nuclear condensates containing Epac1 and general components of the SUMOylation machinery to promote cellular SUMOylation. Furthermore, genetic knockout of Epac1 obliterates oxidized low-density lipoprotein induced cellular SUMOylation in macrophages, leading to suppression of foam cell formation. These results provide a direct nexus connecting two major cellular stress responses to define a molecular mechanism in which cAMP regulates the dynamics of cellular condensates to modulate protein SUMOylation.

## Introduction

Protein SUMOylation is a highly conserved and dynamic post-translational modification and plays important roles in maintaining cellular homeostasis. SUMOylation regulates numerous cellular processes, including transcription, chromatin organization, DNA repair, macromolecular assembly, and signal transduction (*1*). While SUMOylation has long been associated with stress responses, integrating a diverse array of cellular stress signals that trigger rapid increases in global protein SUMOylation (*2-5*), how these cellular stresses promote SUMOylation remains a mystery. In addition, unlike ubiquitination that is tightly regulated by a large number of ubiquitin processing enzymes, including two E1 ubiquitin-activating enzymes, 30-50 E2 ubiquitin-conjugating enzymes, and over 600 E3 ubiquitin ligases (*6*), protein SUMOylation is controlled by a single pair of SUMO-activating enzyme (AOS1/UBA2) E1/SUMO-conjugating enzyme (UBC9) E2 and a minimal set of validated E3 ligases (*7*). The scarcity of SUMO ligases, which are often not required for SUMOylation, suggests that SUMOylation is chiefly controlled globally at the level of E1/E2. However, a general mechanism for the regulation of cellular SUMOylation is lacking.

The cAMP second messenger is a major stress-response signal found to play important roles in diverse biological functions. In vertebrates, the effects of cAMP are mainly transduced by two ubiquitously-expressed intracellular cAMP receptors, the classic protein kinase A (PKA) and the more recently discovered exchange proteins directly activated by cAMP (Epac1 and Epac2) (*8-10*). Extensive studies, particularly recent *in vivo* analyses of Epac1 functions using genetic knockout mouse models and pharmacological probes, reveal that Epac1 regulates a wide range of physiological and pathophysiological processes in response to cellular stresses (*11-15*). Conversely, the expression of Epac1 is often upregulated to promote pathogeneses in various disease models. For example, recent studies demonstrate that enhanced Epac1 expression stimulates pathogenic angiogenesis through simultaneous activation of VEGF and inhibition of Notch signaling in endothelial cells (*16*) and atherosclerosis via promoting the uptake of oxidized low-density lipoprotein (ox-LDL) in macrophages (*17*). Surprisingly, the relationship between cAMP signaling and protein SUMOylation, two common cellular stress response mechanisms, has not been examined and remain unknown. In this study, we demonstrate that cAMP/Epac1 acts as a regulator of SUMOylation via promoting the formation of nuclear condensates containing Epac1 and components of SUMOylation machinery. This Epac1-regulated cellular SUMOylation is physiological important as deletion of Epac1 blocks ox-LDL induced cellular SUMOylation and foam cell formation.

## Results

### SUMOylation machinery is enriched in Epac1-associated proteome

To explore the cellular functions of Epac1, we performed an unbiased Epac1 associated proteome analysis via affinity purification of Epac1-containing cellular complexes in HeLa cells stably expressing an Epac1-FLAG. Shotgun proteomics analyses led to the identification of ∼497 proteins co-immunoprecipitated with only Epac1-FLAG in the anti-FLAG pull-down fraction but not in the control mock immunoprecipitation of HeLa cells stably transfected with an empty vector. These identified Epac1 associated proteins contain many known Epac1 interacting partners, including Annexin A2 (*18*), Importin β1 (*19, 20*), Nup98 (*20*), RanBP2 (*20, 21*), RanGap1 (*19*) and tubulin (*22*). Functional enrichment analysis revealed that the top three enriched pathways associated with the Epac1 proteome are SUMOylation related (**Figure 1A**). Among these identified Epac1-associated proteins, eighteen were identified to overlap with the 81 entries of the protein SUMOylation Gene Ontology (M13502) set. These proteins encompasses all major components of the SUMOylation machinery, including, SUMO-activating enzyme E1 (AOS1/UBA2), SUMO-conjugating enzyme E2 (UBC9), SUMO1/2/3, RanBP2, and PIAS1 (**Figure 1B**). Hypergeometric probability calculation revealed that the Epac1 associated proteome was highly-enriched with SUMOylation related proteins with a representation (enrichment) factor of 8.4 and p value of 6.4×10^−13^ assuming a total human proteome size of 17,874 (*23*). In addition to major components of SUMOylation machinery, Epac1 associated proteome is also highly enriched with well-known SUMO target proteins such as RanGap1, SART1 and HNRNPs. 52 Epac1 associated proteins were found in a database of 689 SUMO target proteins identified by at least 5 previous SUMO proteomic studies (*24*), representing a 2.7 folds enrichment (p = 7.1×10^−11^). To validate the proteomic analysis and to determine if Epac1 can interact with SUMO-E1 directly, we performed a reverse pull-down using nickel affinity resin loaded with purified His-tagged AOS1/UBA2 as a bait. Indeed, we observed specific co-purification of GST-Epac1, but not GST, along with His-AOS1/UBA2 (**Figure S1)**.

**Figure 1.**
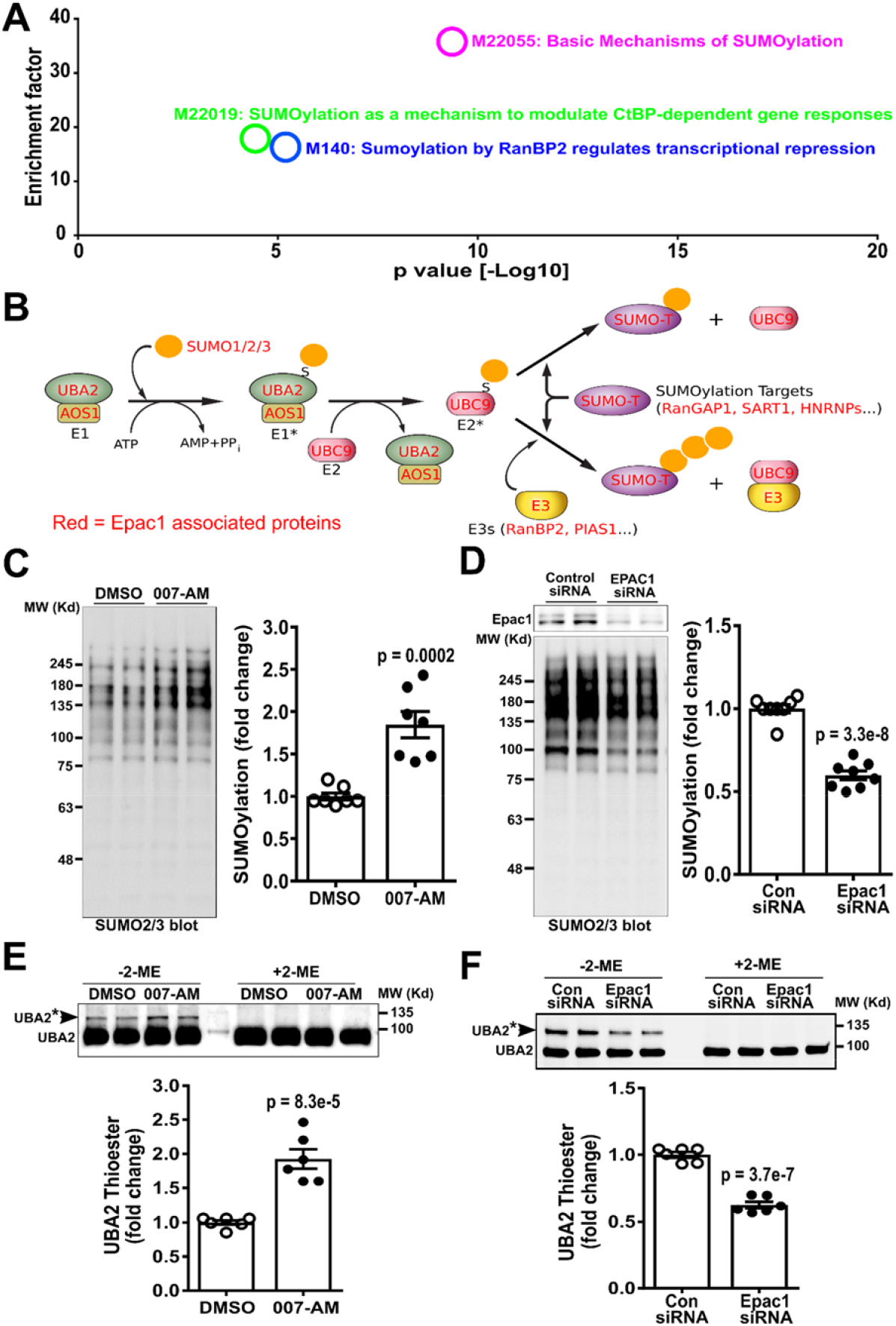
Epac1 associates with SUMOylation machinery and promotes cellular SUMOylation. (A) Top pathways enriched in Epac1 associated proteome revealed by functional enrichment analysis. Basic Mechanism of SUMOylation (M22055) was the most enriched pathway associated Epac1 with an enrichment factor of 35.8 and an associated p value of 4.6×10^−10^. (B) Schematic representation of cellular SUMOylation pathways with proteins found in Epac1 associated proteome highlighted in red. (C) Levels of cellular SUMOylation probed by immunoblotting analysis using anti-SUMO2/3 antibody in HUVECs treated with DMSO or Epac-specific agonist, 007-AM (5 µM) for 30 min. (D) Levels of cellular SUMOylation in HUVECs transfected with control or Epac1-specific siRNA. (E) Levels of UBA2 SUMO-thioester intermediates (UBA2*) examined by immunoblotting analysis using anti-UBA2 antibody in HUVECs treated with DMSO or 5 µM 007-AM (30 min). (F) Levels of UBA2* in HUVECs transfected with control or Epac1-specific siRNA. Data were normalized to total protein loading for SUMOylation or to free UBA2 for UBA2* and shown as Mean ± SEM.

### Epac1 activation promotes cellular SUMOylation

Various cellular stresses such as heat-shock, hypoxia, and osmotic stress trigger cellular SUMOylation (*2-5*). The unexpected finding that Epac1 associated proteome is enriched with the general SUMOylation machinery prompted us to test if activation of Epac1, an effector of a major stress-response signal cAMP, promote cellular SUMOylation. When Human Umbilical Vein Endothelial Cells (HUVECs), which express Epac1 abundantly, but not Epac2 (*18, 25*), were treated with a membrane-permeable Epac-specific agonist, 8-CPT-2’-O-Me-cAMP-AM, also known as 007-AM (*26*), we observed a consistent increase in SUMO2/3-based cellular SUMOylation levels (**Figure 1C**), whereas a control compound PO_4_-AM3 (*27*) had no effect on cellular SUMOylation (**Figure S2A**). On the other hand, SUMO1-based cellular SUMOylation was unchanged in response to 007-AM treatment (**Figure S2B**). Activation of HUVECs with isoproterenol (ISO), a β-adrenergic receptor agonist also led to an enhanced cellular SUMOylation by SUMO2/3 while pretreatment with H89, a PKA specific inhibitor, had no effect on ISO induced cellular SUMOylation (**Figure S2C**). Conversely, suppressing Epac1 by gene silencing using Epac1-specific siRNA led to a reduced total cellular SUMOylation (**Figure 1D**). These results suggest that cAMP acts through Epac1, but not PKA, to promote cellular SUMOylation. 007-AM promoted cellular SUMOylation in HUVECs to a similar extent as induced by heat-shock, which is known to promote global sumoylation (*4*). Furthermore, heat-shock induced global SUMOylation was significantly reduced in HUVEC treated with EPAC1-specific siRNA as compared to HUVEC cells treated with control siRNA (**Figure S3**).

The observed enhancement of cellular SUMOylation by Epac1 activation prompted us to determine if Epac1 activates cellular SUMO E1 and E2 enzymes. When we monitored enzyme activities by probing the level of UBA2-SUMO thioester intermediate (UBA2*) or UBC9-SUMO thioester intermediate (UBC9*) under non-reducing conditions without 2-mercaptoethanol (2-ME) in SDS sample buffer (*28*), activation of Epac1 with 007-AM led to an increase in endogenous cellular UBA2* (**Figure 1E**). However, the majority of cellular UBC9 under the basal unstimulated condition appeared to already existed in the thioester intermediate state in HUVECs and only a slight increase of UBC9* was observed (**Figure S2D**). On the other hand, silencing Epac1 in HUVECs using Epac1-specific siRNA reduced the levels of UBA2* (**Figure 1F**) and UBC9* (**Figure S2E**). Epac1 activation by 007-AM or silencing by siRNA had no effect on free UBA2 level (**Figure S2F&G**).

### Epac1-induced cellular SUMOylation is not dependent on its exchange activity

To determine if Epac1-induced cellular SUMOylation is mediated by its guanine nucleotide exchange (GEF) activity, i.e., its orthodox down-stream effectors Rap1/2, we suppressed cellular Rap1/2 activity by ectopic expression of Rap1GAP, a Rap1/2 specific GTPase-activating protein that efficiently keeps Rap1/2 in their inactive GDP-bound states (*29*). While expression of Rap1GAP in HUVECs blocked the ability of 007-AM to activate Akt, a Rap-dependent Epac1 function (*30*), the basal SUMOylation level (**Figure S4A**) and the ability 007-AM to promote cellular SUMOylation were not affected by Rap1GAP (**Figure S4B**). These results suggest that the effects of Epac1 activation on cellular SUMOylation are not dependent on its canonical down-stream effectors Rap1/2.

### Epac1 does not directly activates SUMO E1 and E2 in vitro

Since Rap1/2 was not required for Epac1-mediated SUMOylation activation and Epac1 interacted with the SUMOylation machinery, we questioned if Epac1 promoted SUMOylation by directly activating SUMO E1/E2. To test this hypothesis, we expressed and purified individual recombinant components of SUMOylation machinery, AOS1/UBA2, UBC9, and SUMOs. We performed *in vitro* SUMO E1/E2 thioester formation experiments. Epac1 was not able to enhance UBC9 thioester formation in the presence or absence of cAMP (**Figure 2**). These results suggest that Epac1 does not activate SUMO E1/E2 directly *in vitro*.

**Figure 2.**
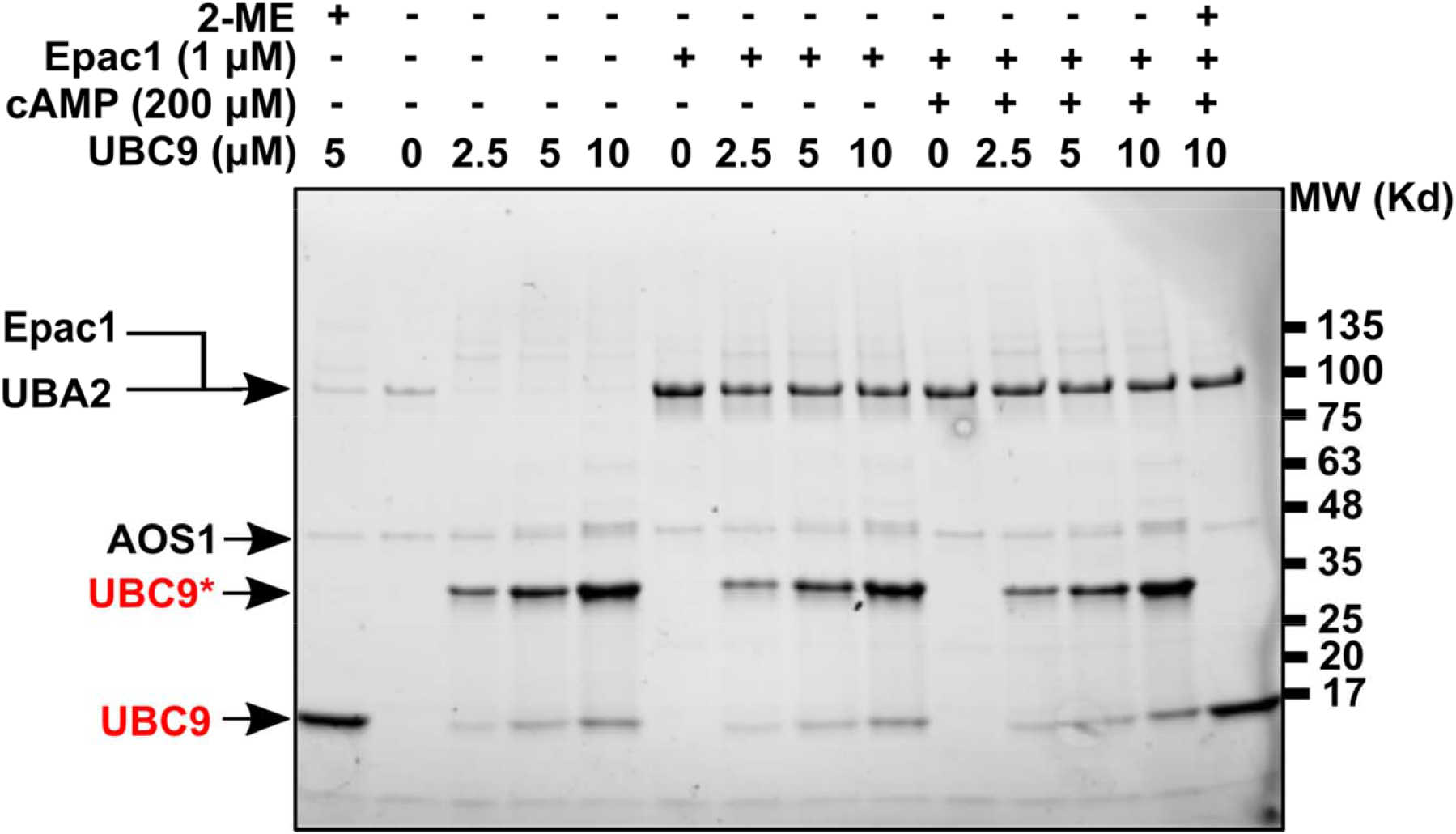
Effects of purified Epac1 protein on UBC9 SUMO thioester formation. In vitro SUMO thioester intermediate formation of recombinant UBC9 in the presence or absence of Epac1 or Epac1 plus cAMP.

### Epac1 activation promotes the formation of Epac1 nuclear condensates that co-localize with nuclear UBA2/UBC9 condensates

The findings that Epac1-induced cellular SUMOylation was not dependent on its GEF activity and that Epac1 did not directly activate SUMO E1/E2 suggest that Epac1 promotes cellular SUMOylation through an unconventional mechanism. We performed confocal microscopy to study the subcellular localization of endogenous Epac1, UBA2, and UBC9 in response to Epac1 activation. Co-immunofluorescence staining of Epac1 and UBA2 and Structured Illumination Microscopy (SIM) super-resolution imaging in HUVECs revealed that under the basal condition, Epac1 staining exhibited distinct cellular puncta both in cytosolic and nuclear compartments (**Figure 3A**). UBA2 labeling displayed a similar puncta staining and distribution with more numerous nuclear puncta than those of Epac1. Significant overlaps between Epac1 and UBA2 signals were observed in nuclear compartments (**Figure 3B**). Activation of Epac1 by 007-AM led to a significant increase in numbers, sizes and intensity of nuclear puncta for Epac1 both in the nuclear and cytosol while the intensity and numbers of nuclear puncta for UBA2 were also modestly increased (**Figure 3A-C**). Moreover, correlation analyses of colocalization using CellProfiler (*31*) revealed that 007-AM stimulation led to a significant enhancement of co-localization of Epac1 and UBA2 nuclear puncta based on both pixel-and object-based approaches (**Figure 3A, D, & E**). Similarly, co-immunofluorescence staining of Epac1 and UBC9 and SIM analysis showed significant overlaps between Epac1 and UBC9 nuclear puncta in HUVECs and treatment with 007-AM increases Epac1 and UBC9 nuclear puncta, as well as their colocalization (**Fig. S5**).

**Figure 3.**
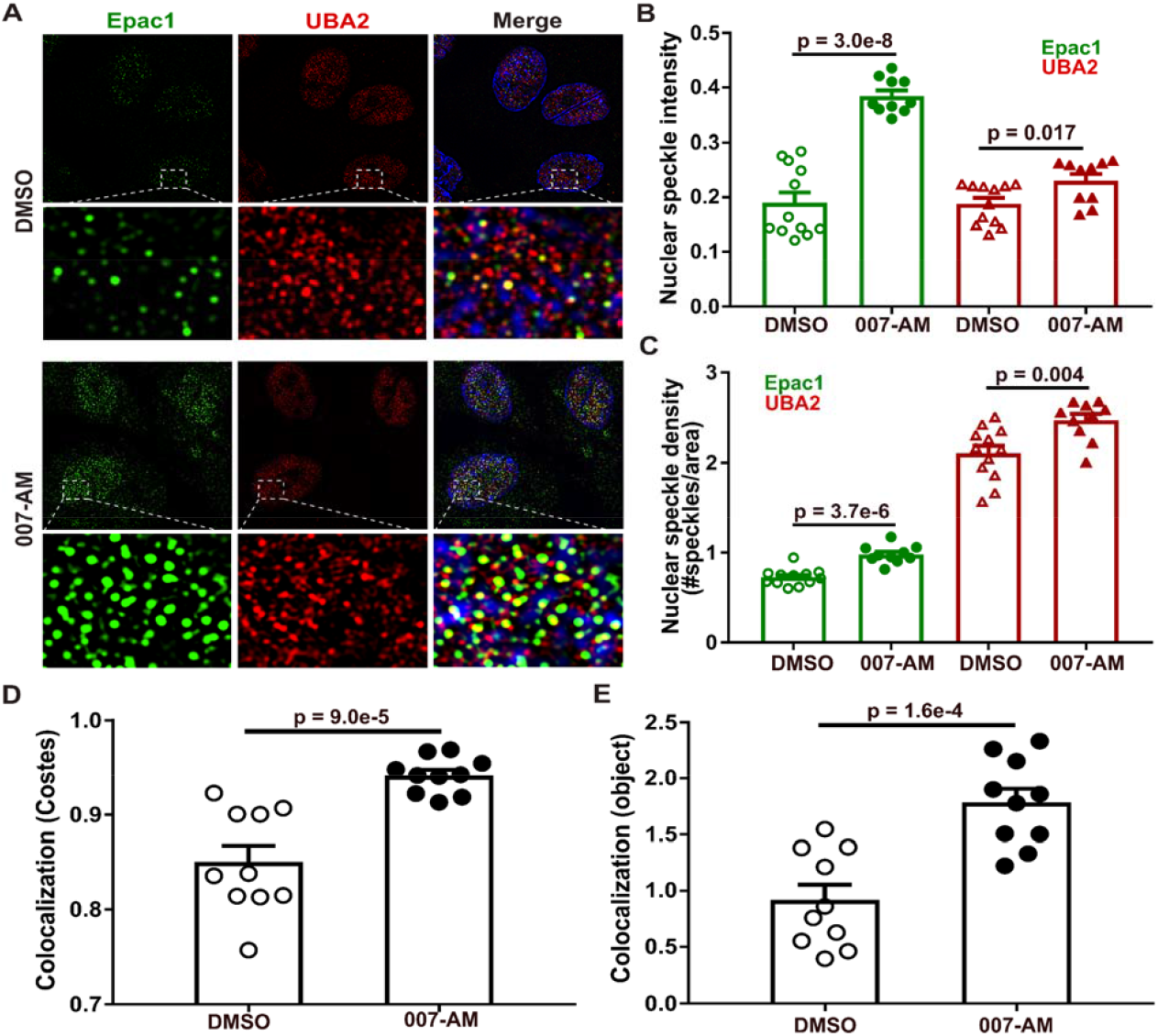
Epac1 activation promotes the formation and colocalization of Epac1 and UBA2 nuclear condensates. (A) SIM immunofluorescence images of endogenous Epac1 (green) and UBA2 (red) probed by anti-Epac1 (SC-25632), and UBA2 (SC-376305) antibodies in control (DMSO) and 007-AM (5 µM, 7 min) treated HUVEC cells. (B) Quantification of Epac1 and UBA2 nuclear speckle intensity in DMSO or 007-AM treated HUVEC cells. (C) Quantification of Epac1 and UBA2 nuclear speckle density in DMSO or 007-AM treated HUVEC cells. (D) Pixel-based colocalization analysis of Epac1 over UBA2 nuclear speckle in DMSO or 007-AM treated HUVEC cells. (E) Object-based colocalization analysis of Epac1 and UBA2 nuclear speckles in DMSO or 007-AM treated HUVEC cells. Images analyses were performed using the CellProfiler software. Data are shown as Mean ± SEM.

### 007-AM and heat-shock induce the formation of Epac1-EYFP/mRuby-UBA2 nuclear condensates

To further characterize the cellular behavior and function of Epac1 and UBA2 cellular condensates, we performed confocal live cell imaging of fluorescently tagged Epac1 and UBA2 in HEK293 cells. Under the basal condition, Epac1-EYFP signals were mostly diffused throughout the cells with enhanced signals concentrated around the nuclear envelope and plasma membrane. A few puncta were observed in the cytosol but mostly absent in the nuclear compartment (**Figure 4A**). These observations are consistent with earlier publications on Epac1-EYFP subcellular localization (*20, 21, 32*). Stimulation of cells with 007-AM led to a robust increase in Epac1-EYFP puncta, particularly in the nuclear compartment (**Figure 4A & B**). In addition, activation of cells with ISO resulted in similar increases in Epac1-EYFP cellular condensates (**Figure S6A**). The induction of Epac1-EYFP nuclear puncta occurred rapidly, almost instantaneously, after the addition of 007-AM (**Figure 4C**). To confirm that the formation of cellular condensates in response to 007-AM is due to direct Epac1 activation, we performed parallel experiments using an Epac1-R279E-EYFP construct that is defective in cAMP binding (*30*). Indeed, mutation of a critical cAMP binding residue abolished the ability of 007-AM or ISO to promote the formation of Epac1-based nuclear condensates (**Figure 4A & B, Figure S6B**). We further performed fluorescence recovery after photobleaching (FRAP) analysis to assess fluidity of 007-AM induced Epac1-EYFP nuclear condensates. The maximal fluorescence intensity of individual Epac1-YFP nuclear condensates recovered quickly after photobleaching, with half-lives ranging from ∼30 to 100s (**Figure S7**).

**Figure 4.**
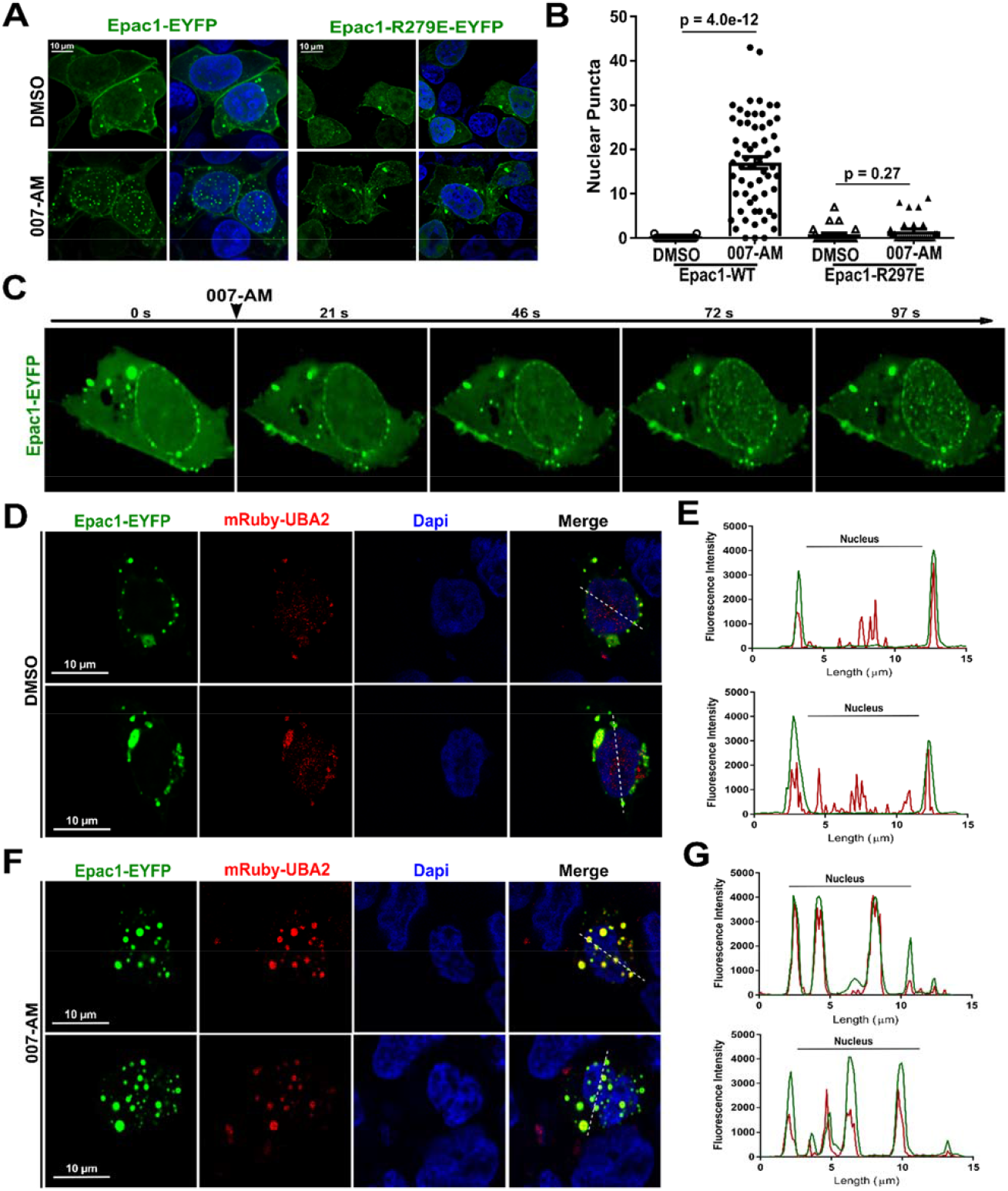
Characterization of Epac1-EYFP and mRuby-UBA2 nuclear condensates. (A) Confocal images of HEK293 cells expressing WT Epac1-EYFP or Epac1-R279E-EYFP in response to 5 µM 007-AM (7 min). (B) Quantification of Epac1-EYFP or Epac1-R279E-EYFP nuclear condensates in DMSO or 007-AM treated HEK293 cells. Data are shown as Mean ± SEM. (C) Confocal live-cell images from a time-lapse movie of Epac1-EYFP expressing HEK293 cells in response to 5 µM 007-AM. (D) Confocal images of HEK293 cells expressing Epac1-EYFP and mRuby-UBA2. (E) Line graphs show fluorescence intensities of Epac1-EYFP and mRuby-UBA2 across the white dashed lines. (F) Confocal images of HEK293 cells expressing Epac1-EYFP and mRuby-UBA2 in response to 5 µM 007-AM. (G) Graphs show fluorescence intensities of Epac1-EYFP and mRuby-UBA2 across the white dashed lines in 007-AM (5 µM) treated HEK293 cells.

When expressed at approximately 30% of the endogenous UBA2 level (**Figure S8**), mRuby-UBA2 showed a similar distribution as endogenous UBA2 staining with diffused speckles mainly in the nuclear compartment and sporadic vacuolar-like structures in the cytosol that partially overlapped with the Epac1-EYFP signals, particularly around the nuclear envelope under unstimulated basal condition (**Figure 4D & E**). Strikingly, in response to 007-AM stimulation, mRuby-UBA2 speckles coalesced to form larger nuclear condensates that were superimposable with nuclear Epac1-EYFP puncta (**Figure 4F & G**). Since both 007-AM and heat-shock promoted cellular SUMOylation, we asked if heat-shock could also induce nuclear Epac1-EYFP and mRuby-UBA2 condensates as 007-AM. Indeed, when HEK293 cells expressing Epac1-EYFP and mRuby-UBA2 were subjected to heat-shock at 43 °C, a time-dependent formation of co-localized Epac1-EYFP and mRuby-UBA2 nuclear condensates were observed (**Figure 5**). These results suggest that 007-AM and heat-shock induced cellular SUMOylation potentially shares a similar mechanism.

**Figure 5.**
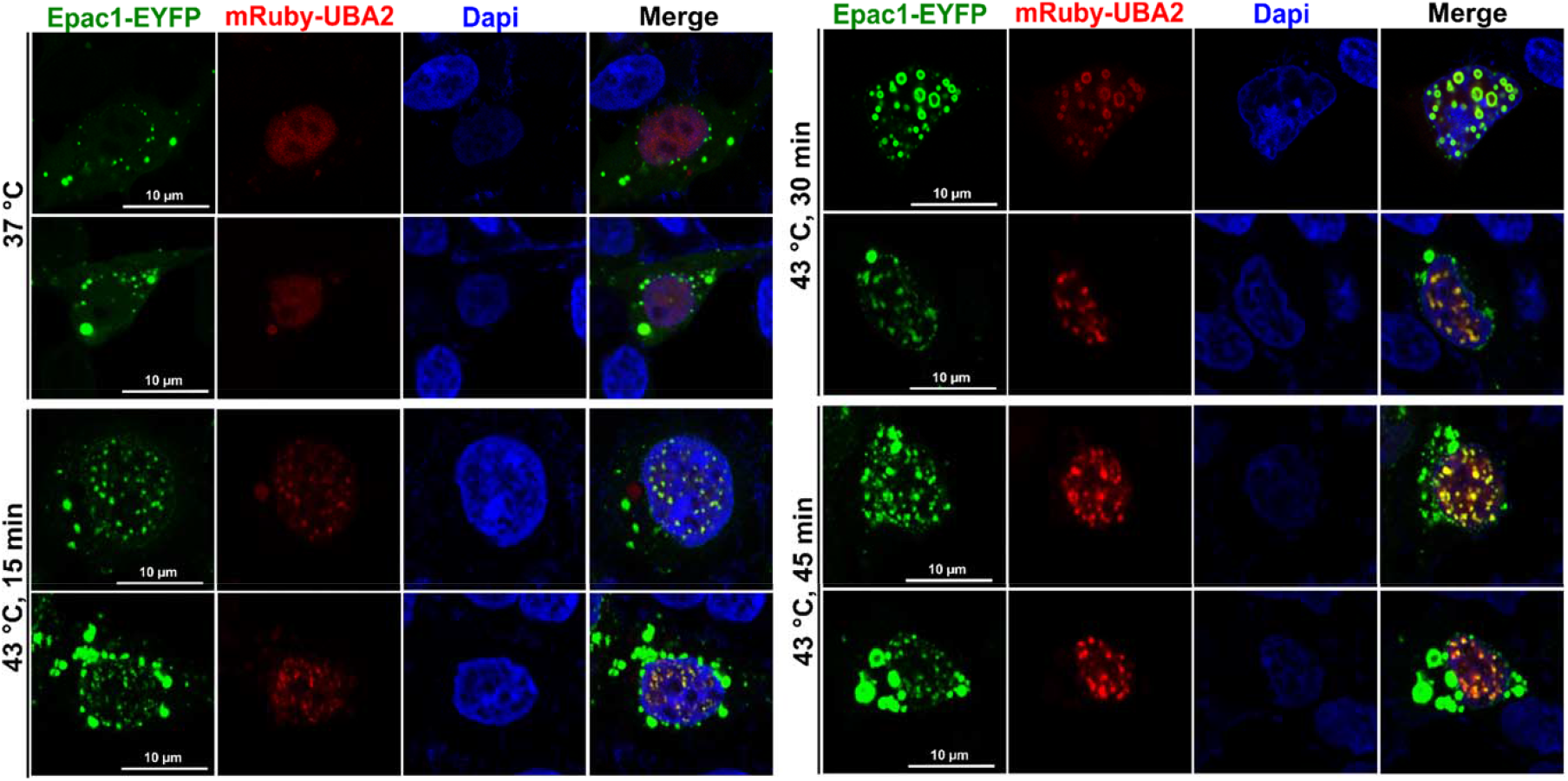
Heat-shock induces the formation of colocalized Epac1-EYFP and mRuby-UBA2 nuclear condensates in HEK293 cells. Confocal images of HEK293 cells expressing Epac1-EYFP and mRuby-UBA2 at 37 °C or 43 °C for 15, 30 and 45 min

Numerous membrane-less nuclear condensates, such as nuclear speckles, PML (promyelocytic leukaemia) bodies, PcG (Polycomb Group) bodies, etc. are known to exist and involved in a wide array of important cellular functions (*33*). The discovery of cAMP-dependent formation of Epac1 nuclear condensates prompted us to investigate whether these Epac1 nuclear puncta conformed to a previously described nuclear bodies. When 007-AM treated Epac1-EYFP expressing HEK293 cells were stained for various nuclear body markers, including Serine/arginine-rich splicing factor (SC35), PML protein, and Polycomb complex protein (BMI-1), the numbers of Epac1 nuclear condensates were much higher than those of the known nuclear bodies. In addition, no significant overlap was observed between Epac1-EYFP puncta and nuclear speckles or PcG bodies while a small numbers of Epac1 nuclear condensates partially co-localized with the PML bodies (**Figure S9**). These results suggest that Epac1 nuclear puncta likely define a new type of nuclear condensate structure.

### Epac1 contains intrinsically disorder regions and undergoes cAMP-dependent phase separation (PS)

The discovery of Epac1-based cellular condensates suggests the Epac1 belongs to a growing family of proteins capable of modulating biological functions via undergoing PS. One of the common feature shared by many proteins prone to PS is the presence of intrinsically disorder regions (IDRs) with multiple interacting motifs (*34*). Sequence analysis of Epac1 protein by IUPred (*35*) and PONDR (*36*) indeed revealed multiple potential IDRs (**Figure 6A**). This notion is in agreement with the fact that purified recombinant Epac1 protein requires high salt (500 mM) to maintain solubility, and that Epac1 is recalcitrant to crystallization and readily undergoes PS in the presence of various polyethylene glycols when subjected to crystallization screenings (**Figure 6B**). Furthermore, when diluted to low salt concentrations (150 mM), Epac1 protein solution underwent reversible aggregation and became cloudy. On the other hand, in the presence of 3.5% 1,6 hexanediol, low salt Epac1 solution remained clear (**Figure 6C**). 1,6 hexanediol is a PS indicator used to trigger the dissolution of liquid-like assemblies but not solid-like aggregations (*37-39*). To further characterize the PS properties of Epac1 and the effect of cAMP, we determined the saturation concentration (C_sat_) of Epac1 as a function of salt concentrations in the presence or absence of cAMP via light scattering measurements. In the absence of cAMP, increasing salt concentrations increased the C_sat_ (**Figure 6D**) as expected. Noticeably, addition of cAMP dramatically decreased C_sat_ across all ionic strength conditions (**Figure 6E**), suggesting that cAMP promotes Epac1 phase separation.

**Figure 6.**
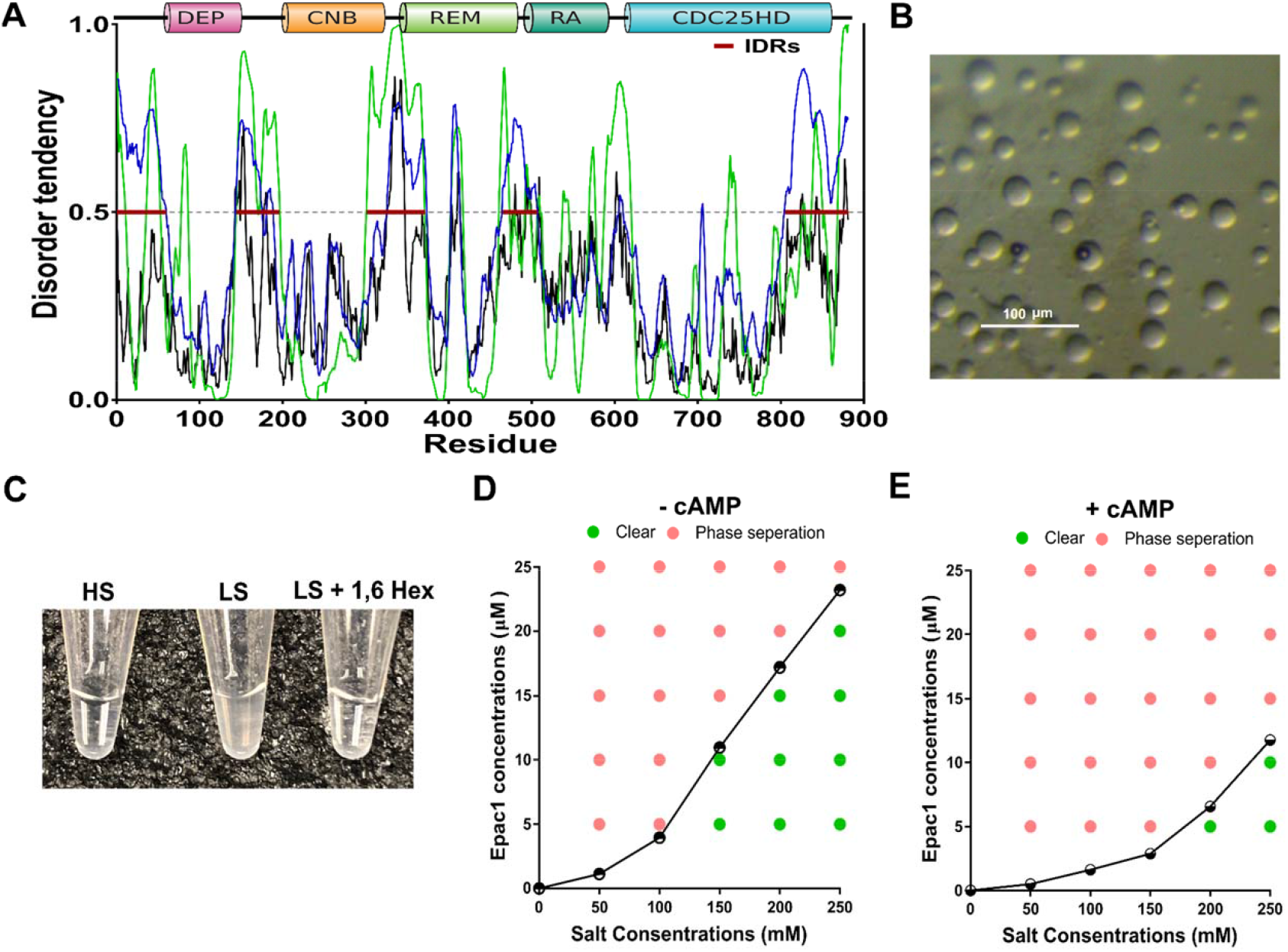
Regulation of Epac1 PS by ionic strength and cAMP. (A) Disorder tendency scores of Epac1 predicted by IUPred (black) and PONDR (green: VLXT score; blue: VLS2 score). (B) Microscopic images of liquid-liquid PS of Epac1 (2.5 mg/ml) in 50 mM Sodium acetate pH 4.5, 100 mM Lithium sulfate, 25 % w/v PEG 400. (C) 15 μM recombinant purified Epac1 protein in the presence of 500 mM (HS), 150 mM (LS) NaCl, and LS with 3.5% 1,6 hexanediol (1, 6 Hex). (D) Epac1 phase diagrams showing C_sat_ as a function of salt concentrations. (E) Epac1 phase diagrams in the presence of 30 μM of cAMP.

### Epac1 condensates are required for the Epac1-mediated activation of SUMOylation

Having demonstrated that Epac1, UBA2 and UBC9 colocalize and form nuclear condensates, we next asked if the formation of Epac1 nuclear condensates are required for Epac1-mediated cellular SUMOylation. To address this question, we screened various Epac1 mutants and identified a previously well-characterized Epac1 deletion construct, Δ(1-148)Epac1 that has the first 148 amino acids deleted at the N-terminus (*40-43*). This mutant has increased solubility/stability when expressed recombinantly but retains all measurable biochemical properties such as cAMP binding and Rap activation (*40, 41*). Moreover, GST-Δ(1-148)Epac1 was capable of interacting with AOS1/UBA2 as the full-length GST-Epac1 (**Figure 7A**). On the other hand, unlike full-length Epac1, purified Δ(1-148)Epac1 was soluble at all protein and salt concentrations tested and did not undergo PS in the presence or absence of cAMP. Consistent with this notion, when Δ(1-148)Epac1-YFP was expressed in HEK293 cells, unlike Epac1-EYFP, this construct failed to form nuclear condensates in response to 007-AM stimulation (**Figure 7B**). Therefore, this deletion mutant is ideal for testing if the ability of Epac1 to form nuclear condensate is important for promoting cellular SUMOylation. When Δ(1-148)Epac1 was expressed in HUVEC cells, 007-AM was no longer able to promote cellular SUMOylation (**Figure 7C**) and UBA2 thioester bond formation (**Figure 7D**). Taken together, these results suggests that cAMP-mediated Epac1 activation promotes the formation of Epac1 nuclear condensates, which are responsible for cAMP-induced cellular SUMOylation and UBA2 activation.

**Figure 7.**
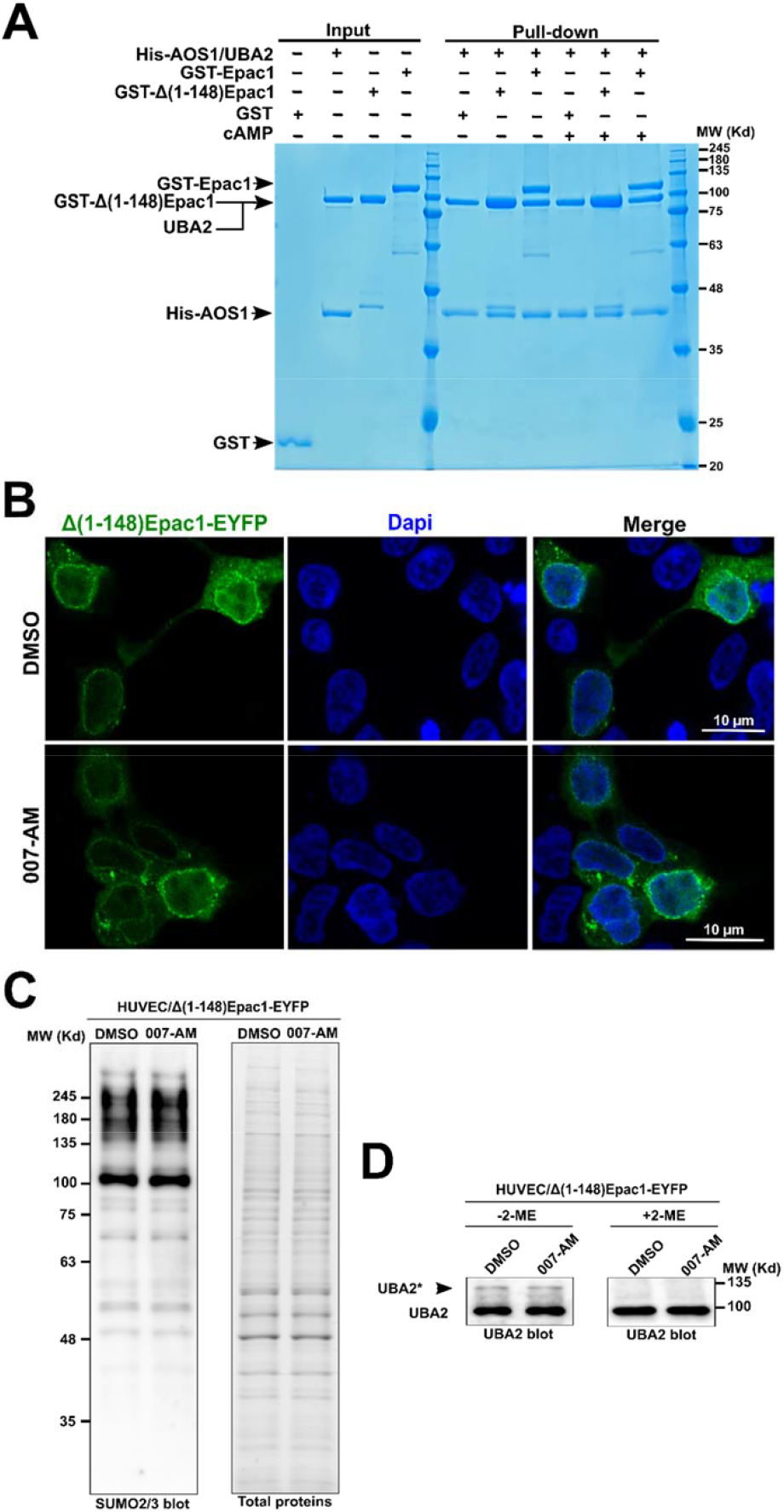
Formation of nuclear Epac1 condensates is required for the Epac1-mediated activation of SUMOylation. (A) Affinity pull-down of recombinant purified GST-Δ(1-148)Epac1 and GST-Epac1 by His-AOS1/UBA2 in the presence or absence of cAMP (50 µM). (B) Confocal images of HEK293 cells expressing Δ(1-148)Epac1-EYFP in response to vehicle or 007-AM (5 µM, 7 min) treatment. (C) Levels of cellular SUMOylation probed by immunoblotting analysis using anti-SUMO2/3 in Δ(1-148)Epac1-EYFP expressing HUVECs treated with DMSO or 007-AM (5 µM, 7 min). (D) Levels of UBA2 SUMO-thioester intermediates (UBA2*) examined by immunoblotting analysis using anti-UBA2 antibody in Δ(1-148)Epac1-EYFP expressing HUVECs treated with DMSO or 007-AM (5 µM, 7 min).

### Epac1 is required for ox-LDL stimulated cellular SUMOylation and foam cell formation

Encouraged by our findings that Epac1 activation promoted SUMOylation in cells, we further asked if Epac1 plays a role in regulating SUMOylation under physiological settings. Our recent studies demonstrated that deletion of Epac1 in an atherogenic mouse model reduced atherosclerotic plaque formation by suppressing ox-LDL mediated foam cell formation (*17*). Surprisingly, when we isolated bone marrow derived monocytes (BMDMs) from wild-type (WT) and Epac1-knockout mice and challenged them with ox-LDL, we observed an increase in cellular SUMOylation by SUMO2/3 in the WT BMDMs (**Figure 8A**). Our recent study has shown that ox-LDL can induce intracellular cAMP (*17*), pointing to the possibility that ox-LDL induced SUMOylation may be mediated in part by cAMP and associated down-stream effectors. Indeed, ox-LDL mediated increase in cellular SUMOylation was subdued in Epac1 null BMDMs (**Figure 8A**), suggesting that Epac1 is responsible for ox-LDL mediated increases in cellular SUMOylation. Next, we tested if Epac1 activation alone was sufficient to promote cellular SUMOylation and UBA2 activation in BMDMs. Stimulation of BMDMs by 007-AM led to a significant increase in cellular SUMOylation in WT BMDMs, but not in Epac1 null BMDMs, in a similar manner to the response to ox-LDL stimulation (**Figure 8B**). Consistent with increased cellular SUMOylation, UBA2-SUMO thioester intermediate levels were concomitantly increased in WT BMDMs following ox-LDL treatment. On the other hand, the UBA2-SUMO thioester levels in Epac1 null BMDMs were not affected by ox-LDL treatment (**Figure 8C**). Moreover, when cells were treated with 007-AM, we observed a significant increase in UBA2-SUMO thioester levels in the WT BMDMs, but not in Epac1-null BMDMs (**Figure 8D**). Taken together, these results demonstrate that Epac1 activation is sufficient and necessary to induce ox-LDL-mediated cellular SUMOylation and UBA2 activation in primary BMDMs.

**Figure 8.**
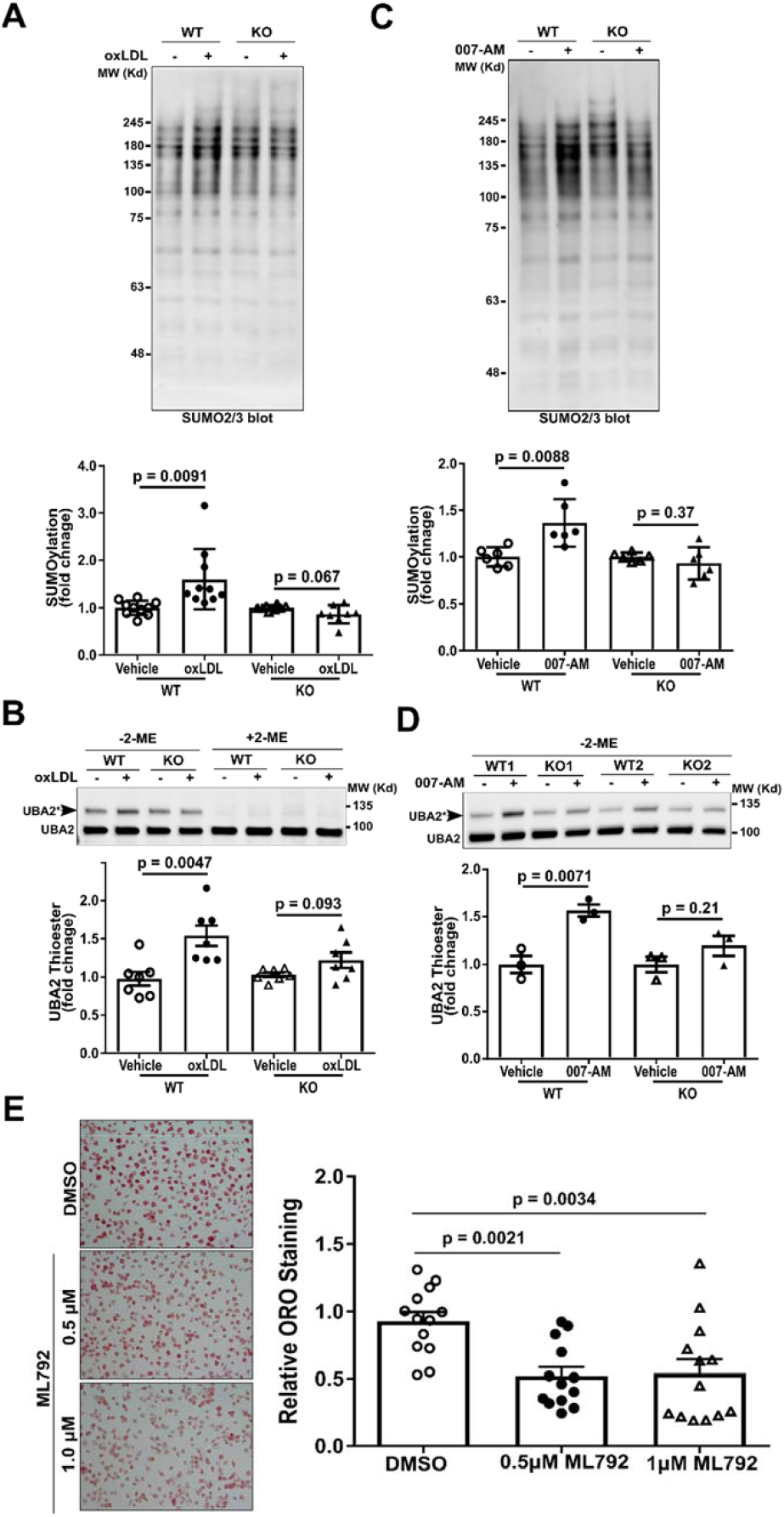
ox-LDL and 007-AM promote cellular SUMOylation and UBA2 activation in an Epac1-dependent manner. Cellular SUMOylation probed by anti-SUMO2/3 antibody in WT and Epac1^-/-^ (KO) BMDMs treated with vehicle or 40 µg/mL ox-LDL for 7 min (A) or with vehicle or 5 µM 007-AM for 30 min (B). Formation of UBA2-SUMO thioester (UBA2*) in WT and Epac1^-/-^ BMDMs in response ox-LDL (C) or 007-AM stimulation (D). (E) Representative images and quantification of Oil-Red-O stained mouse primary macrophages treated with 40 µg/mL ox-LDL in the presence or absence of UBA2 inhibitor ML792. Data are shown as Mean ± SEM.

Our previous studies have shown that ox-LDL increases intracellular cAMP and upregulates Epac1 expression in macrophages supporting the importance of Epac1 for ox-LDL mediated foam cell formation (*17*). To determine if cellular SUMOylation is involved in this process, we pretreated the BMDMs with ML792(*44*). As expected, treatment of BMDMs with ML792 suppressed cellular SUMOylation (**Figure S10**). Importantly, ML792 significantly blocked ox-LDL uptake in macrophages (**Figure 8E**), suggesting that cellular SUMOylation is functionally important for ox-LDL uptake in macrophage and foam cell formation. Collectively, our studies reveal a previous unknown mechanism, in which ox-LDL acts through cAMP/Epac1 to promote SUMOylation-dependent foam cell formation.

## Discussion

Protein SUMOylation has long been associated with stress responses. However, a molecular mechanism linking cellular stresses and SUMOylation is missing. Our studies reveal that Epac1 is associated with the general SUMOylation machinery and that cAMP, an ancient and universal stress-response second messenger, acts through Epac1 to regulate cellular SUMOylation. Unexpectedly, Epac1 does not act through its canonical effectors, Rap1/2 GTPase, to promote cellular SUMOylation. Instead, enhanced SUMOylation by Epac1 activation is accompanied by the formation of nuclear condensates containing Epac1 and components of the general SUMOylation machinery. These results provide a direct nexus connecting two major cellular stress responses and define a molecular mechanism in which cAMP controls the dynamics of Epac1 cellular condensates to promote protein SUMOylation.

While protein SUMOylation has been implicated to promote phase separation (PS) by enhancing weak multivalent interactions between SUMO and SUMO interacting motif within various protein binding partners (*45*), our findings provide a novel mechanism in which cAMP/Epac1 regulates SUMOylation through promoting the formation of cellular condensation. The ability of LLPS/biomolecular condensates to enhance catalysis has been well-documented (*46*). A recent study reports that SUMOylation rates are significantly enhanced in an artificial system where the SUMOylation machinery is recruited into engineered condensates generated by PS of multidomain scaffolding proteins (*47*). These results provide theoretical supports for our findings that the rapid formation of Epac1 SUMO-activating nuclear condensates can act as SUMOylation organizers where concentrated SUMOylation machinery and substrates accelerate cellular protein SUMOylation via mass action and/or substrate channeling in response to cellular stresses or environmental signals. While we could not rule out that Epac1 only regulates a subset of SUMO targets in a cell-type specific manner, our data as a whole support the concept that Epac1 condensates act as “SUMOylation organizers” to generally enhance SUMOylation, particularly considering that heat-shock, a well-known trigger for global SUMOylation, induced the formation of Epac1/UBA2 nuclear condensates and that silencing Epac1 attenuated heat-shock induced global SUMOylation.

The observation of cAMP-dependent Epac1 PS correlates very well with the cellular behaviors of Epac1 condensates. Under basal low cAMP conditions when Epac1 exists in the apo conformation with higher C_sat_, the number and size of the Epac1 cellular condensates are smaller than those of stimulated conditions where activation of Epac1 by cAMP leads to a decreased C_sat_ that promotes the formation of Epac1 cellular condensates (**Figure 9**). These observations of cAMP-induced Epac1 condensates, coupled with a recent report of cAMP-modulated PKA regulatory subunit RI PS (*48*), suggest that cAMP is an important molecular switch/trigger for biomolecular condensate regulation. The ability of cAMP to directly modulate the dynamics of biomolecular condensates provides the first experimental evidence that protein phase separation can be regulated by an endogenous ligand, and opens up a new dimension in our understanding of this ancient stress response second messenger.

**Figure 9.**
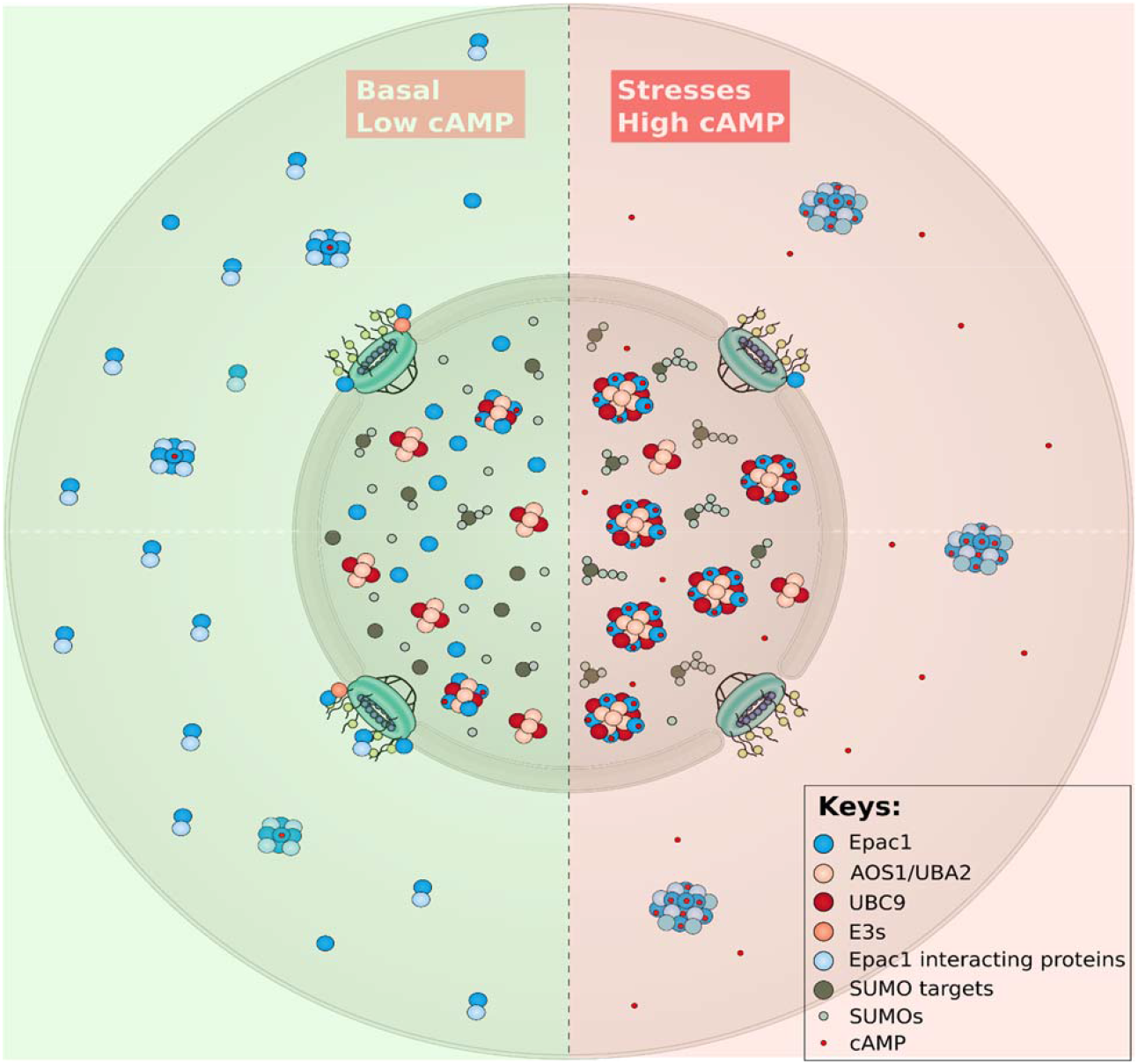
Regulation of Epac1 cellular condensates by cAMP. Schematic model of Epac1-mediated cellular condensates under basal (low cAMP) and stress (high cAMP) conditions.

In summary, we discover that cAMP, acting through Epac1, promotes the formation of SUMO-activating nuclear condensates and enhances cellular SUMOylation. These findings represent a major conceptual advance in our understanding of cellular stress responses by providing a direct connection between protein SUMOylation and cAMP signaling, two major cellular stress processes. Considering the universal presence of protein SUMOylation and cAMP signaling in biology, our studies have major implications for various physiological and pathophysiological functions, such as the cardiovascular and neuronal systems where both cAMP/Epac1 signaling and protein SUMOylation play significant roles (*14, 49-52*). Since dysregulations of Epac1 signaling have been implicated in the development of numerous pathophysiological conditions, including cancer (*53, 54*), chronic pain (*55-57*), infections (*58, 59*), and vascular proliferative diseases (*16, 17, 60-62*), it is possible that hyperactivation of Epac1 might induce pathogenesis in part through activating cellular SUMOylation. Furthermore, the ability of cAMP/Epac1 signaling, a highly coordinated and compartmentalized process, to directly regulate the dynamics of PS suggests that the formation of biomolecular condensates represents another important mechanism for organizing cellular space and functions in addition to the classic membrane-based intracellular compartmentalization.

## Materials and Methods

### Reagents

Dulbecco’s Modified Eagle’s Medium (DMEM) - high glucose (Cat. # D5796), Fetal bovine serum (FBS, Cat. # F2442), Isoproterenol hydrochloride (Cat. # I5627), C_12_E_9_, PROTEIN GRADE^®^ Detergent, 10% Solution (Cat. #205534), Glutathione Sepharose^®^ 4B Fast Flow (Cat. #GE17-0756), N-Ethylmaleimide (NEM, Cat. # E3876), Ni Sepharose® Fast Flow (Cat. # GE17-5318), Poly-L-Lysine Solution (0.01%, Cat. # P4707) and Gelatin solution (Cat. # G1393-100ML) were from MilliporeSigma. 8-(4-Chlorophenylthio)-2’-O-methyladenosine-3’,5’-cyclic monophosphate acetoxymethyl ester (007-AM) was from Axxora (Cat. # BLG-C051). DAPI (Cat. # 62248), Hoechst 33342 Solution (Cat. #62249) and Lipofectamine^®^ 2000 (Cat. # 11668-019) were from ThermoFisher Scientific. cOmplete^™^, Mini, EDTA-free Protease Inhibitor Cocktail Tablet was from Roche (Cat. # 11836170001). Protein A/G PLUS-Agarose (Cat. # sc-2003) was from Santa Cruz Biotechnology, Inc. FluorSave™ Reagent (Cat. # 345789) was from EMD Millipore. Tris(2-carboxyethyl)phosphine hydrochloride (TCEP, Cat. # HR2-651) was from Hampton Research. Recombinant mouse macrophage colony stimulating factor (rm-MCSF) was from R&D System (Cat. # 416-ML). Human highly oxidized LDL was obtained from KB Kalen Biomedical (Cat. # 770252-6). ML792 (Cat. # 407886) was purchased from MEDKOO.

### Cell culture and transfection

Human umbilical vein endothelial cells (Lonza, Cat. # C2519A) were maintained and sub-cultured in EGM-2™ Endothelial Cell Growth Medium (Lanzo, Cat. # CC-3162) at 37°C in a 5% CO_2_ humidified incubator. Cells passages between 2 and 8 were used for experiments described in this study. For experiments involving RNA interference (RNAi), HUVECs at 70% confluence were transfected with Epac1-specific (ThermoFisher Scientific, Cat. # 1299001) or non-targeting control Stealth RNAiTM siRNA oligonucleotides (ThermoFisher Scientific, Cat. # 12935300) at a final concentration of 50 nM. HEK-293 cells (ATCC, Cat. # CRL-1573) were maintained in DMEM with high glucose supplemented with 10% FBS. Cell transfection was performed using Lipofectamine® 2000 according to the manufacturer’s instructions.

### Epac1 associated proteome analyses by shotgun proteomics

To identify potential Epac1 cellular binding partners, we performed immunoaffinity purification of cellular Epac1-containing protein complexes from HeLa cells stably expressing full-length Epac1 with a C-terminal FLAG/HA tandem epitope tag as previously described (*18, 22*). Briefly, Epac1-FLAG/HA from two 10 cm plates of cell lysates was captured by immunoprecipitation using anti-FLAG antibody-conjugated agarose beads. The bound protein complexes were eluted with purified FLAG peptide. Control mock purification was performed using HeLa cells stably transfected with an empty vector to excluding nonspecific interactions. The immunoprecipitation eluents were loaded onto 10% SDS-PAGE gel. Shortly after all the eluents were migrated into the gels, gel bands (1-2 cm) containing the total protein loading were excised and subjected to in-gel tryptic digestion following a well-established protocol as described previously (*63*). The tryptic digested samples were analyzed by LC/MS/MS using an Orbitrap Fusion™Tribrid™mass spectrometer (Thermo Scientific™) interfaced with a Dionex UltiMate 3000 Binary RSLCnano HPLC System. One microgram of each sample was loaded for the analysis. Peptides were separated with an Acclaim™ PepMap™ C_18_ column (75µm ID x 15 cm) at a flow rate of 300 nl/min. Gradient conditions were: 3%-22% B for 40 min, 22%-35% B for 10min, 35%-90% B for 10 min, 90% B held for 10 min, (Buffer A, 0.1 % formic acid in water; Buffer B, 0.1% formic acid in acetonitrile). The peptides were analyzed using the data-dependent acquisition (DDA) mode. The survey scan was performed with 120K resolution at 400 *m/z* from 350 to 1500 *m/z* with AGC target of 2e^5^ and max injection time of 50 msec. The DDA cycle was limited to 3 seconds. Monoisotopic masses were then selected for further fragmentation for ions with 2 to 5 plus charge within a dynamic exclusion range of 35 seconds. Fragmentation priority was given to the most intense ions. Monoisotopic precursor selection was enabled. Precursor ions were isolated using the quadrupole with an isolation window of 1.6 m/z. CID was applied with a normalized collision energy of 35% and resulting fragments were detected using the rapid scan rate in the ion trap. The AGC target for MS/MS was set to 1e^4^ and the maximum injection time was limited to 35 msec. The raw data files were processed using Thermo Scientific™ Proteome Discoverer™ software version 1.4. The spectra were searched against the Uniprot Homo sapiens database using SEQUEST search engine. The database search was restricted with the following parameters. Trypsin was set as the enzyme with maximum missed cleavages set to 2. The precursor ion tolerance was set to 10 ppm, and the fragment ion tolerance was set to 0.8 Da. Variable modifications were set to methionine oxidation and phosphorylation on serine, threonine and tyrosine. Carbamidomethylation on cysteine was set as a static modification. The search results were validated and trimmed to a 5% FDR using Percolator. Proteins identified only in Epac1-FLAG pull down but not in the mock control were designated as Epac1-associated proteome. Functional enrichment analysis was performed using the ToppGene Suite (*64*). Enrichment factor was computed by hypergeometric probability calculation. To visualize highly enriched pathways (MSigDB and KEGG), we plotted resulting enrichment factor against the p-values.

### Cellular SUMOylation and UBA2/UBC9-SUMO thioester formation assays

Near confluent HUVECs or BMDMs in 12-well plate were washed once with Hank’s Buffered Salt Solution (HBSS). HUVECs were starved in serum-free (SF) Endothelial Basal Medium (EBM) and then treated with DMSO, PO4-AM3 1.67 µM), 007-AM (5 µM), ISO (20 µM), H89 (5 µM), ISO plus H89, or heat shock at 43 °C for 30 min or otherwise indicated. BMDMs were first serum deprived in DMEM High Glucose with 2.5% FBS, 1× Non-Essential Amino Acids and 100 μg/ml Penicillin/Streptomycin for 2 hours, followed with two quick rinses with SF DMEM and resuspended in 400 μl of SF DMEM for treatment. After 5 min to allow cells to adjust to SF conditions, control vehicles, 007-AM (5µM) or ox-LDL (40 µg/mL) were added to cells for 30 min or 7 min, respectively. After treatment, cells were washed twice with warm DPBS and lysed with 100 µl 1×SDS sample buffer (50 mM Tris, pH 6.8, 2% SDS, 0.1% Bromophenol blue, 3% 2-mercaptoethanol (2-ME) and 10% glycerol) with protease inhibitors and 20 mM NEM. Total cell lysates were collected and sonicated on ice using 15W power output for three to four cycles of 5 s with 5 s rests in between until complete soluble. After heat denaturation at 95 °C for 5 min, the samples were subjected to immunoblotting analysis of cellular SUMOylation using an anti-SUMO2/3 (Enzo Life Sciences, Inc., Cat.# MBL-PW9465) or anti-SUMO1 antibody (Enzo Life Sciences, Inc., Cat.# MBL-PW8330). For the UBA2 or UBC9 thioester bond formation assay, the cell lysates were collected in SDS sample buffer without 2-ME and split into two equivalent volumes, then 2-ME was added to one of the two samples before loading onto SDS-PAGE gels for immunoblotting analyses with anti-UBA2 (Abcam, Cat. # ab185955) or anti-UBC9 antibody (Abcam, Cat. # ab185955).

### Immunoblotting analysis

Protein samples from cultured cells or immunoprecipitation were resolved on stain-free SDS-PAGE gels. After electrophoresis, images were captured by ChemiDoc™ Touch Imaging System (Bio-Rad) for total protein loading quantification before proteins were transferred to PVDF membrane (Millipore, Cat. # IPVH00010). The blots were incubated with primary antibodies against SUMO2/3 (Enzo Life Sciences, Cat. # BML-PW9465 or MBL International, Cat. # M114-3), Epac1 (Cell Signaling Technology,, Cat. # 4155), UBA2 (Abcam, Cat. # ab185955), UBC9 (Abcam, Cat. # ab185955), FLAG (MilliporeSigma, Cat. # F1804), PML (Santa Cruz Biotechnology, Cat. # SC966), and TRIM28 (ProteinTech, Cat. # 66630) at 4 °C for overnight followed by incubation with horseradish peroxidase-conjugated secondary antibodies (Bio-Rad) and detection using Amersham™ ECL™ Prime Western Blotting Detection Reagent (GE Healthcare Life Sciences, Cat. #45-002-401). The chemiluminescence signals were captured with a ChemiDoc™ Touch Imaging System (Bio-Rad) and quantitated using Image Lab™ Software (Bio-Rad) or ImageJ software. Individual signal of a specific protein band was first normalized against corresponding total protein loading and the final immunoblotting readout was expressed as a ratio of the normalized treatment signal to the normalized control signal. Statistical analysis was performed using data from at least three independent experiments.

### Recombinant protein expression and purification

Recombinant full-length human Epac1 and Δ(1-148)Epac1 proteins were constructed as a Glutathione S-transferase (GST)-fusion, expressed in *Escherichia coli* CK600K cells and purified as described previously (*43, 65*). Epac1 and Δ(1-148)Epac1 without the GST-tag was generated by thrombin cleavage of GST-Epac1and further purified to more than 95% purify using a Superdex 200 FPLC column. Recombinant His_6_-AOS1/UBA2, UBC9, SUMO1 and SUMO2/3 proteins were expressed and purified as described previously (*66*).

### Affinity pull-down of GST-Epac1 or GST-Δ(1-148)Epac1 by His_6_-AOS1/UBA2 recombinant proteins

0.1 nanomole purified His_6_-AOS/UBA2 protein (>90% purity) was immobilized on Ni Sepharose beads (30 ul of 50% slurry per sample) in a loading buffer (50 mM Tris, pH 7.5, 150 mM NaCl, 1 mM MgCl_2_, 1 mM EDTA, 0.5% C_12_E_9_, 1 mM TCEP, 1× protease inhibitor) at 4 °C with constant gentle mixing for one hour. The beads were then washed and equilibrated in binding buffer (50 mM Tris, pH 8.5, 150 mM NaCl, 1 mM MgCl_2_, 1 mM EDTA, 0.5% C_12_E_9_, 1 mM TCEP, 1× protease inhibitor). 0.1 nanomole purified GST, GST-(1-148)Epac1 or GST-Epac1 protein was added to each reaction mixture in the absence or presence of 50 μM cAMP, respectively. After 45 minutes incubation at 37 °C with constant gentle mixing, the beads was washed twice by 500 μl binding buffer and three times by a washing buffer (same as the binding buffer except with 0.05% C_12_E_9_). His_6_-AOS/UBA2 was eluted from the beads with 30 μl washing buffer B containing 300 mM imidazole and analyzed using SDS-PAGE. 50 μM cAMP was included in all the binding and washing steps for samples contain cAMP.

### In vitro UBC9 thioester intermediate formation assay

UBC9 SUMO-thioester bond formation assay was performed using purified recombinant proteins in a SUMOylation reaction buffer containing 20 mM HEPES, pH 7.3, 110 mM KOAC, 2 mM Mg(OAC)_2_, 1 mM EGTA, 1 mM DTT, and 0.05% Tween 20. All reactions contained 500 nM AOS1/UBA2, 20 μM SUMO2, and varying UBC9 concentrations at 0, 2.5, 5 and 10 μM. If present, Epac1 or cAMP concentration was at 1 μM or 20 μM, respectively. The reaction was initiated with the additional of 2 mM ATP and carried out at 37 °C for 20 min. At the end of reaction, the assay was split into two equal portions, and mixed with 2×SDS sample buffer with or without 2-ME, respectively. After heating at 95 °C for 5 min, the samples were analyzed by electrophoresis using stain-free SDS-PAGE.

### Immunofluorescence staining

HUVECs, HEK293 or BMDMs plated on glass coverslips coated with 2% gelatin or Poly-L-Lysine (10 μg/ml) were treated with 007-AM or vehicle control for 7 min and washed with PBS for two to three times. The cells were fixed with 10% formalin for 15 min at 37 °C, rinsed three times with PBS, 5 min each and permeabilized with 0.25% Triton X-100 for 10 min. After rinsing with PBS for three times, the cells were incubated with 5% normal goat or horse serum in PBS for 30 min to block non-specific binding and followed by incubation of anti-Epac1 (1:50; Santa Cruz Biotechnology, Inc. Cat. # SC-25632 or SC-28366), anti-UBA2 (1:250; Santa Cruz Biotechnology, Inc. Cat. # SC-376305), anti-UBC9 (1:50; Cell Signaling Technology, Cat. # 4786), anti-SUMO1 (1:100; Cell Signaling Technology, Cat. # SC-4930), anti-SUMO2/3 (1:200; Cell Signaling Technology, Cat. # SC-4971), anti-SC35 (1:200; Abcam, Cat. # 11826), anti-PML (1:200; Santa Cruz Biotechnology, Cat. # SC966), anti-BMI (1:500; Cell Signaling Technology, Cat. # 6964) at 4 °C for overnight. After washing with 1×TBST (20 mM Tris, 137 mM NaCl, 0.1% Tween 20) for three times, cell specimens were incubated with goat anti-Rabbit IgG antibody, DyLight® 488 (1:200; Vector Laboratories, Burlingame, CA, Cat. #DI-1488) or horse anti-Mouse IgG antibody, DyLight® 549 (1:200; Vector Laboratories, Burlingame, CA, Cat. #DI-2594) for 30 min at room temperature in dark. After washing with 1× TBST for three times, cell nuclei were stained with DAPI solution. Coverslips were mounted with FluorSave™ reagent for fluorescence microscopic imaging.

### SIM fluorescence microscopic imaging and analysis

Images of double immunofluorescence staining of endogenous Epac1 and UBA2 or Epac1 and UNC9 in HUVECs were captured by Nikon n-SIM Structured Illumination Super Resolution Microscope System using 100× oil objective in slice 3D-SIM mode. 405 nm, 488 nm, and 561 nm lasers were used for three-color imaging. All images from both the 007-AM treatment and DMSO control groups were captured using same parameters including the laser power and exposure time. Images for more than eight randomly selected fields from at least three independent coverslips per treatment condition were collected. SIM images were reconstructed using the Open Thumbnail N-SIM Slice Reconstruction Window of NIS-Elements Software (N-SIM) with the same optimized parameters for reconstruction and LUT’s adjustment for all images from both the 007-AM treatment and control groups. The signal intensity and density of the Epac1 and UBA2 nuclear speckles were analyzed using CellProfiler (*31*) with the speckle counting and scoring modules while correlation between Epac1 and UBA2 nuclear speckles was analyzed with the colocalization module.

### Confocal fluorescence microscopic imaging and analysis

Fluorescence images were captured by a Nikon A1R Confocal Laser Microscope System using a 100 × oil objective. Three-dimensional rendering of the fluorescent intensity topography was conducted by using the 3D surface plot analysis in ImageJ software. All images were subject to the same parameters for generation of the contoured images. Four to eight randomly selected fields from at least three independent coverslips per treatment were used for data analysis. The nuclear residing puncta were analyzed in ImageJ (v1.53c) by separating the channels, thresholding the DAPI channel and using particle analysis to automate adding nuclear regions to the Region of Interest (ROI) manager. These ROIs were then applied to the green or red channel where a second thresholding was conducted for high intensity puncta which was held constant across images. Individual nuclear ROIs were selected, watershed process was applied, and particle analysis was used to determine the number of puncta present within a given nucleus. This analysis was applied to a minimum of 24 independent cells from multiple fields of view for each condition. Co-occurrence of fluorescence signal was determined by direct line analysis measurement in ImageJ. Channels for an image were segregated using the split function, followed by applying a straight line through the cytoplasm and nucleus of each cell on one channel and creating an identical line on the second channel using the restore selection function. The plot profile for each image was then taken as list of mean grey intensities or relative fluorescence intensities to generate colored line graphs.

### Live cell imaging

For live cell imaging, HEK293 cells were plated in glass-bottom plates (Mat-Tek Cat. # P35G-1.5-14-C) and transfected with Epac1-EYFP, Epac1-R279E-EYFP, pcDNA-mRuby-UBA2 or combination of Epac1-EYFP and pcDNA-mRuby-UBA2. 24-36 hours after transfection and immediately before live cell imaging, the cells were rinsed with warmed DPBS, incubated in 100 μl phenol-red free DMEM with Hochest nuclear staining dye and treated with 100 μl of vehicle control, 007-AM (5 µM), or ISO (10 µM) in DPBS for live cell imaging with a Nikon A1R confocal microscope. During imaging, cells were placed in a pre-warmed humid chamber heated to 37°C with 5% CO_2._ NIS-elements software was used to set time-lapse capture of a single field of view every second for 10 minutes after addition of 007-AM or ISO. Static confocal images of the same field were captured prior to and after the treatment with agonist as references.

### Fluorescence recovery after photobleaching (FRAP) analysis in live cell

FRAP experiments were conducted using Nikon A1R confocal microscope equipped with a Tokai Hit stage top incubation chamber heated to 37 °C and circulated with 5% CO_2_ air for live cell imaging. Briefly, HEK293 cells seeded in poly-l-lysine coated glass-bottom slide chamber were transfected with Epac1-EYFP. 24 hours after transfection, the cells were starved with phenol-red free DMEM for 1 hour and treated with 5 μM of 007-AM. Nuclear Epac1-EYFP condensates formation was monitored by live cell imaging. Appropriate field with multiple stable Epac1-EYFP nuclear condensates was selected. Baseline fluorescence intensities were established by collecting several frames of images before photobleaching using a 488 nm laser set at 100% power output. Fluorescence recovery was followed by time lapse video capturing at one frame per 10 sec for 210 sec. NIS-Elements Software was used to analyze the fluorescence recovery by time measurement of the maximum pixel intensity of individual condensate within the selected ROI area. Fluorescence intensity was normalized to the baseline fluorescence intensity.

### Disorder tendency and net charge calculation

The disorder tendency was predicted by IUPred2A (*35*) and PONDR (*67*). To plot the sliding net charge, the net charge of a sliding window of 20 residues was computed by assigning residues D and E a charge of -1, K and R a charge of +1, and H a charge of +0.5.

### Epac1 phase separation (PS) diagram

Purified recombinant Epac1 (50 μM) in a HS buffer (25 mM Tris pH 8.8, 500 mM NaCl, 5 mM MgCl_2_ and 1 mM TCEP) was diluted to 5 (1:10), 10 (1:5), 15 (3:10), 20 (4:10) and 25 (1:1) μM concentration using a dilution buffer (same as the HS buffer but with no NaCl), resulting corresponding NaCl concentrations of 50, 100, 150, 200 and 250 mM, respectively, with or without 30 μM final cAMP. The protein solutions were incubated at room temperature for 15 min and the degree of the protein PS was quantified by monitoring the absorbance (light scattering) at 320 nm using a NanoDrop spectrophotometer. To determine the C_sat_, the protein solutions were cleared by centrifugation at 16,000 rpm for 15 min at room temperature and Epac1 concentrations in the supernatant were determined by OD_280_ monitored via wavelength scanning between 240-320 nm using a NanoDrop spectrophotometer. The Epac1 phase diagrams in the presence or absence of cAMP was generated by plotting the Epac1 C_sat_ as a function of salt concentrations (*34*).

### Isolation of primary BMDMs

Four-month-old mice were euthanized by isoflurane followed by cervical dislocation. Femur and tibiae were removed and flushed with 10 ml of Dulbecco’s phosphate-buffered saline (DPBS) using a 25-guage needle to harvest the bone marrow. Resulting cells were siphoned through a 40 µm nylon cell strainer, centrifuged at 500 g for 5 minutes at 4 °C, decanted, and resuspended in red blood cell lysing buffer Hybri-Max for 5 minutes RT. Cells were pelleted and washed twice, then resuspended in 5 ml of FBS to be counted and seeded at 5-6×10^6^ cells per 10 cm uncoated, sterile culture dish for 12-16 h in M_0_ media (DMEM High Glucose, 10% FBS, 20 ng/mL rm-MCSF, 100 μg/ml Penicillin/Streptomycin, 1× Non-Essential Amino Acids) at 37 °C, 5% CO_2_. Floating and loosely adherent cells were transferred to a cell culture-coated well for downstream experiments. After 24 h incubation, an equivalent volume of fresh M_0_ media was added to each dish and incubated for 72 h. Half of the media was replaced with fresh M_0_ media for an additional 48 h then this half-media exchange was repeated, and cells incubated for 24 h to complete the differentiation to naïve macrophages (M_0_) cells before further treatment.

### BMDM differentiation

*Ex vivo* differentiation of M_0_ BMDMs to foam cells (M_foam_) differentiation was accomplished by incubating the M_0_ cells with M_foam_ differentiation media (DMEM High Glucose, 2% FBS, 100 μg/ml Penicillin/Streptomycin, 1× Non-Essential Amino Acids, and 40 µg/mL ox-LDL) for 48 h. Cells were pretreated with either ML-792, at indicated concentrations, or DMSO for an hour prior to addition of the ox-LDL. After differentiation, cells were washed thrice with cold DPBS, and fixed in 10% neutral-buffered formalin for 30 min at 4 °C followed by two additional DPBS washes, and then equilibrated for 5 min in 78% methanol at room temperature. Cells were then stained for 15 min with fresh 0.2% (w/v) Oil Red O (ORO) solution under constant agitation and afterwards de-stained for 1 min with 78% methanol followed by extensive DPBS washes. Following ORO staining, cells were counterstained with Meyer’s hematoxylin and 10 random fields of view were captured for each well on a Nikon light microscope to attain an average visual representation of overall lipid staining. The ORO stain was then eluted with 100% methanol for 10 min for quantitative measure of each well. Eluent was transferred to a 96-well plate where absorbance at 500 nm was measured by a FlexStation® 3 microplate reader (Molecular Devices).

### Statistical analyses

All data were analyzed using the Student’s t-test or the analysis of variance with Bonferroni correction for multiple comparisons between groups. Measurements are expressed as Means ± SEM. Statistical significance was designated as P< 0.05.The Materials and Methods section should provide sufficient information to allow replication of the results. Begin with a section titled Experimental Design describing the objectives and design of the study as well as prespecified components.

## Acknowledgments

We thank Drs. Olga Chumakova and Travis Moore for support and assistance in microscopic imaging.

## Author contributions

Wenli Yang: Methodology; Validation; Formal analysis; Investigation; Data Curation; Writing-Review & Editing; Visualization. William Robichaux: Methodology; Validation; Formal analysis; Investigation; Data Curation; Writing-Review & Editing; Visualization. Fang Mei: Methodology; Validation; Formal analysis; Investigation; Data Curation; Writing-Review & Editing; Visualization. Wei Lin: Methodology; Validation; Investigation; Data Curation; Writing-Review & Editing. Li Li: Methodology; Validation; Investigation; Data Curation. Sheng Pan: Data Curation; Writing-Review & Editing. Mark White: Methodology; investigation. Yuan Chen: Resources; Writing-Review & Editing. Xiaodong Cheng: Conceptualization; Methodology; Validation; Formal analysis; Investigation; Resources; Data Curation; Writing-Original draft preparation, Review & Editing; Visualization; Supervision; Project administration; Funding acquisition.

## Data and materials availability

All data needed to evaluate the conclusions in the paper are present in the paper and/or the Supplementary Materials.

