## Supplemental figures for "Epac1 regulates cellular SUMOylation by promoting the formation of SUMO-activating nuclear condensates"

**Figure S1.** Epac1 interacts directly with SUMO E1.
**Figure S2.** Epac1 activation promotes cellular SUMOylation.
**Figure S3.** Role of EPAC1 in heat-shock-induced cellular SUMOylation.
**Figure S4.** Epac1-mediated cellular SUMOylation is not dependent on its canonical effectors, Rap small GTPases.
**Figure S5.** Epac1 activation promotes the formation of Epac1 and UBC9 nuclear condensates and their colocalization.
**Figure S6.** ISO promotes the formation of cAMP-dependent Epac1 nuclear condensates.
**Figure S7.** FRAP analysis of Epac1-EYFP nuclear condensates.
**Figure S8.** Expression levels of endogenous UBA2 and ectopic mRuby-UBA2 in HEK293 cells.
**Figure S9.** Co-localization analyses of Epac1-EYFP nuclear condensates and common nuclear condensate markers.
**Figure S10.** UBA2 specific inhibitor ML792 blocks cellular SUMOylation in BMDMs.

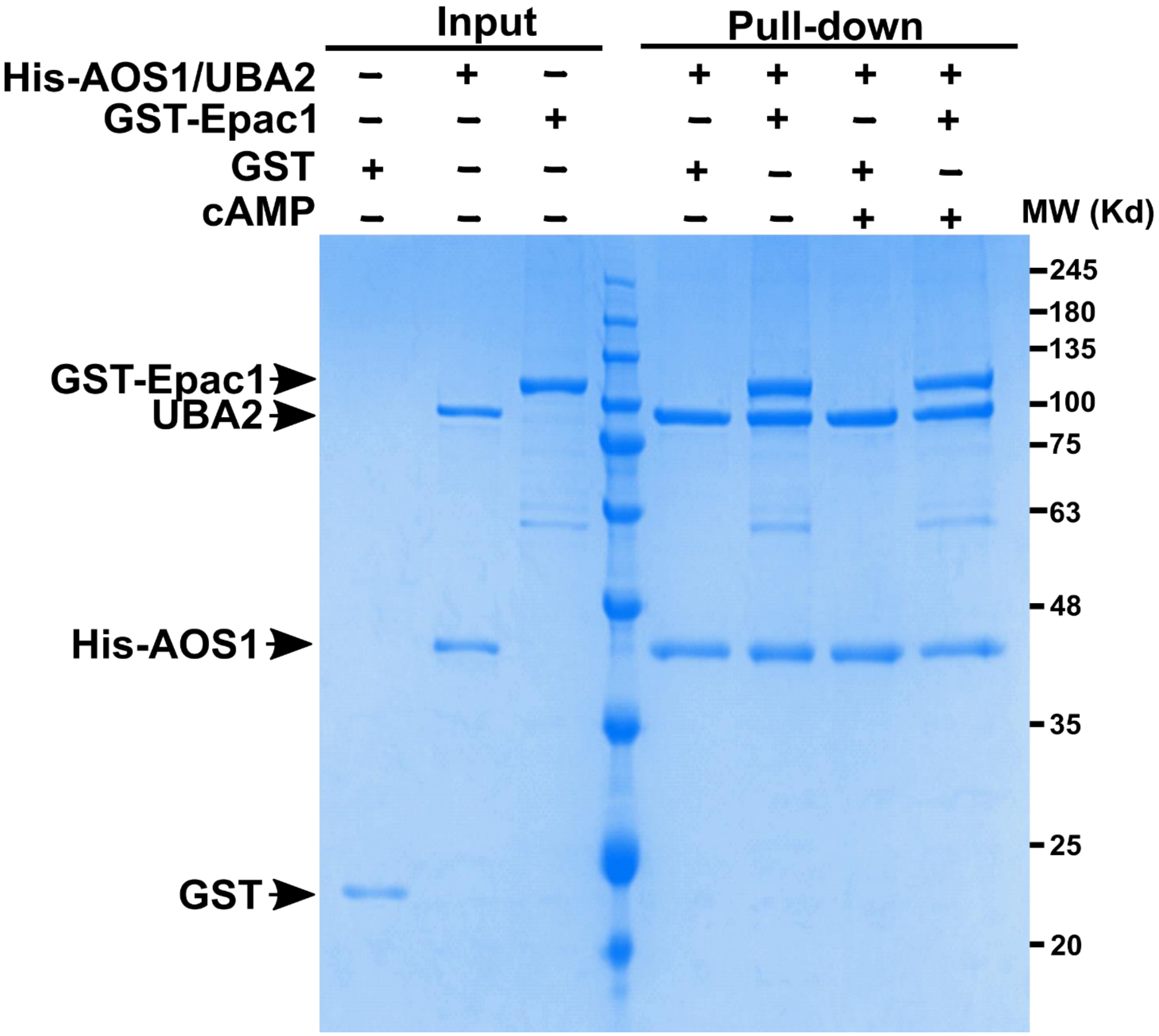


**Figure S1. Epac1 interacts directly with SUMO E1.** Affinity pull-down of purified recombinant GST-Epac1 by His-AOS1/UBA2 in the presence or absence of cAMP (50 μM).

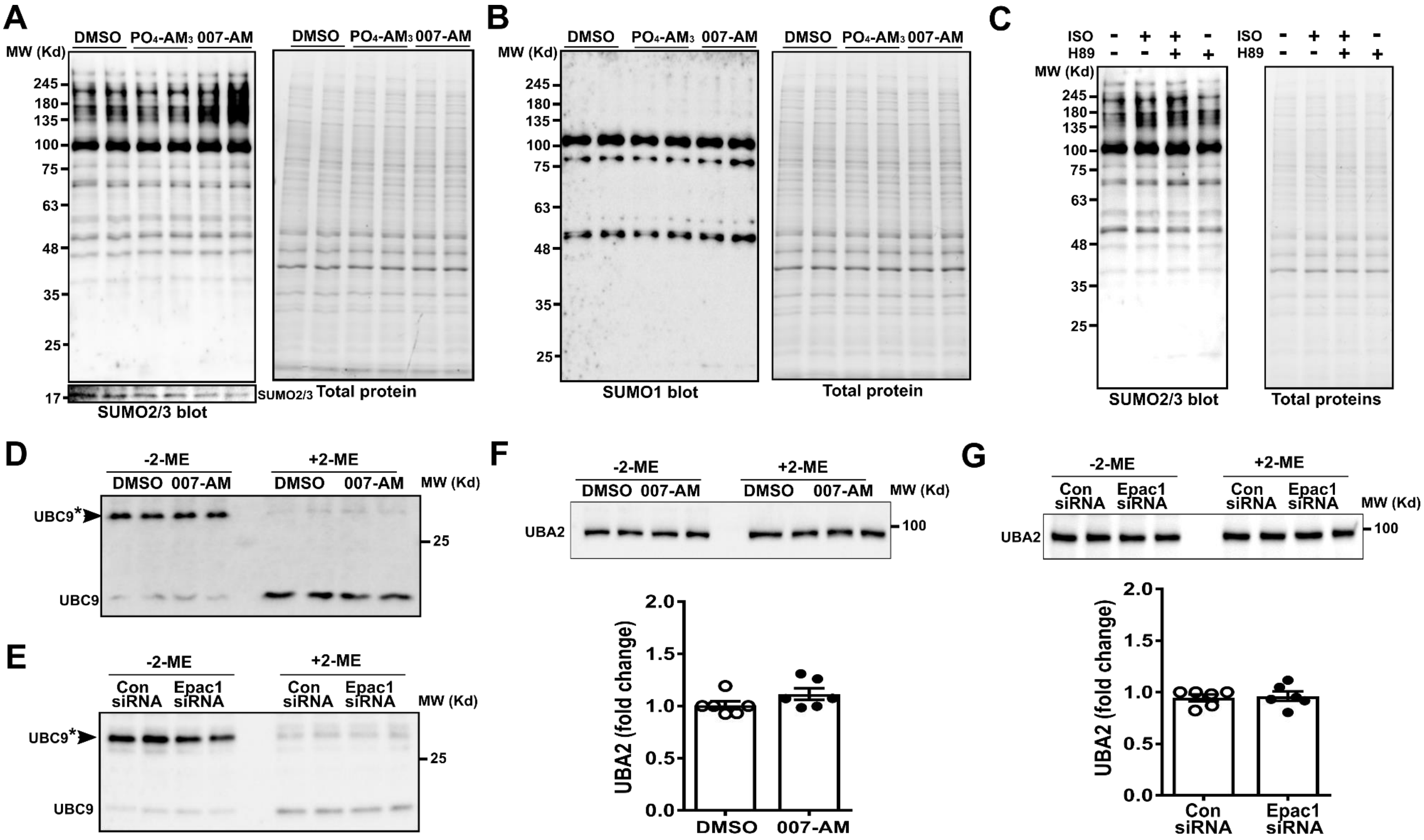


**Figure S2. Epac1 activation promotes cellular SUMOylation.** Levels of cellular SUMOylation probed by immunoblotting analysis using anti-SUMO2/3 (A) or anti-SUMO1 (B) antibody in HUVECs treated with DMSO, PO4-AM3 (1.67 µM) or Epac-specific agonist, 007-AM (5 µM) for 30 min. (C) Levels of cellular SUMOylation probed by anti-SUMO2/3 antibody in HUVECs treated with ISO (20 µM) with or without H89 (5 µM) for 30 min. (D) Levels of UBC9 SUMO-thioester intermediates (UBC9*) examined by immunoblotting analysis using anti-UBC9 antibody in HUVECs treated with DMSO or 5 µM 007-AM (30 min). (E) Levels of UBC9* in HUVECs transfected with control or Epac1-specific siRNA. Quantification of free UBA2 in response to 007-AM (F) or Epac1 siRNA (G). Date were normalized to total protein loading and shown as Mean ± SEM.

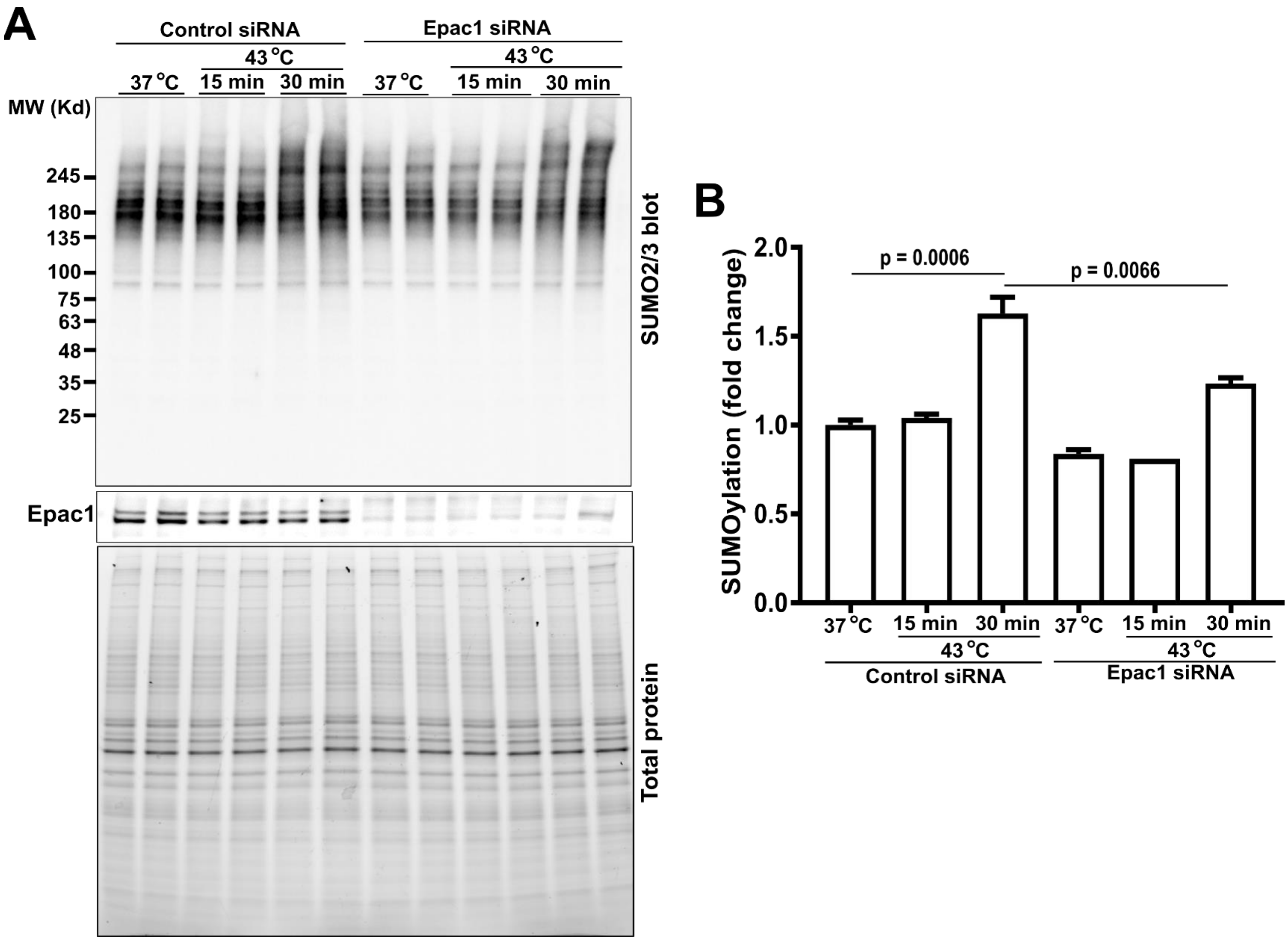


**Figure S3. Role of EPAC1 in heat-shock-induced cellular SUMOylation.** (A) Levels of cellular SUMOylation in HUVECs transfected with control or Epac1-specific siRNA in response to heat-shock treatment (43 °C, 15 or 30 min). (B) Quantification of heat-shock induced cellular SUMOylation in HUVECs. Date were normalized to total protein loading and shown as Mean ± SEM (N = 4).

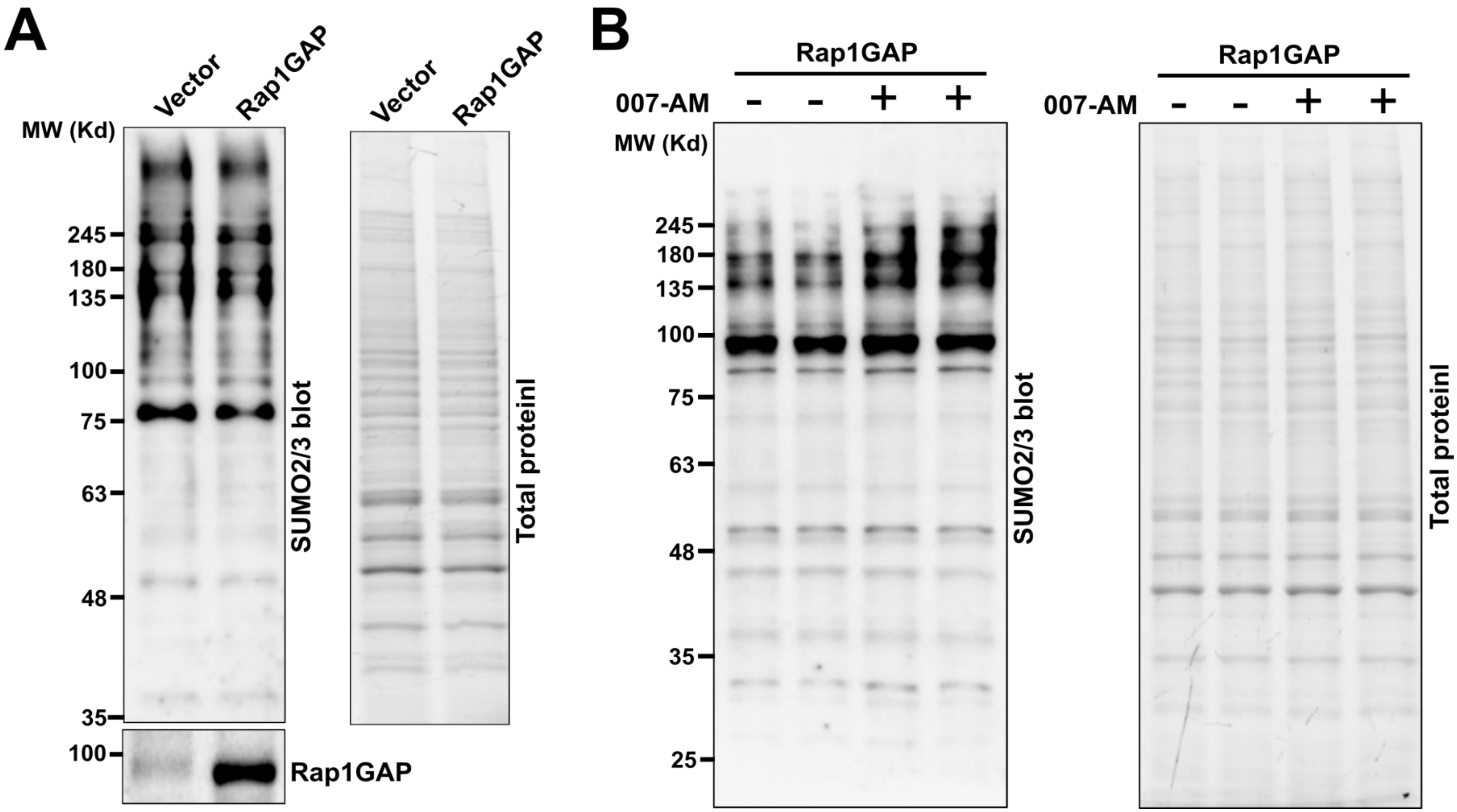


**Figure S4. Epac1-mediated cellular SUMOylation is not dependent on its canonical effectors, Rap small GTPases.** Levels of cellular SUMOylation probed by immunoblotting analysis using anti-SUMO2/3 antibody in HUVECs transfected with Rap1GAP or empty vector (A), and in response to DMSO or Epac-specific agonist, 007-AM (5 μM) for 30 min (B).

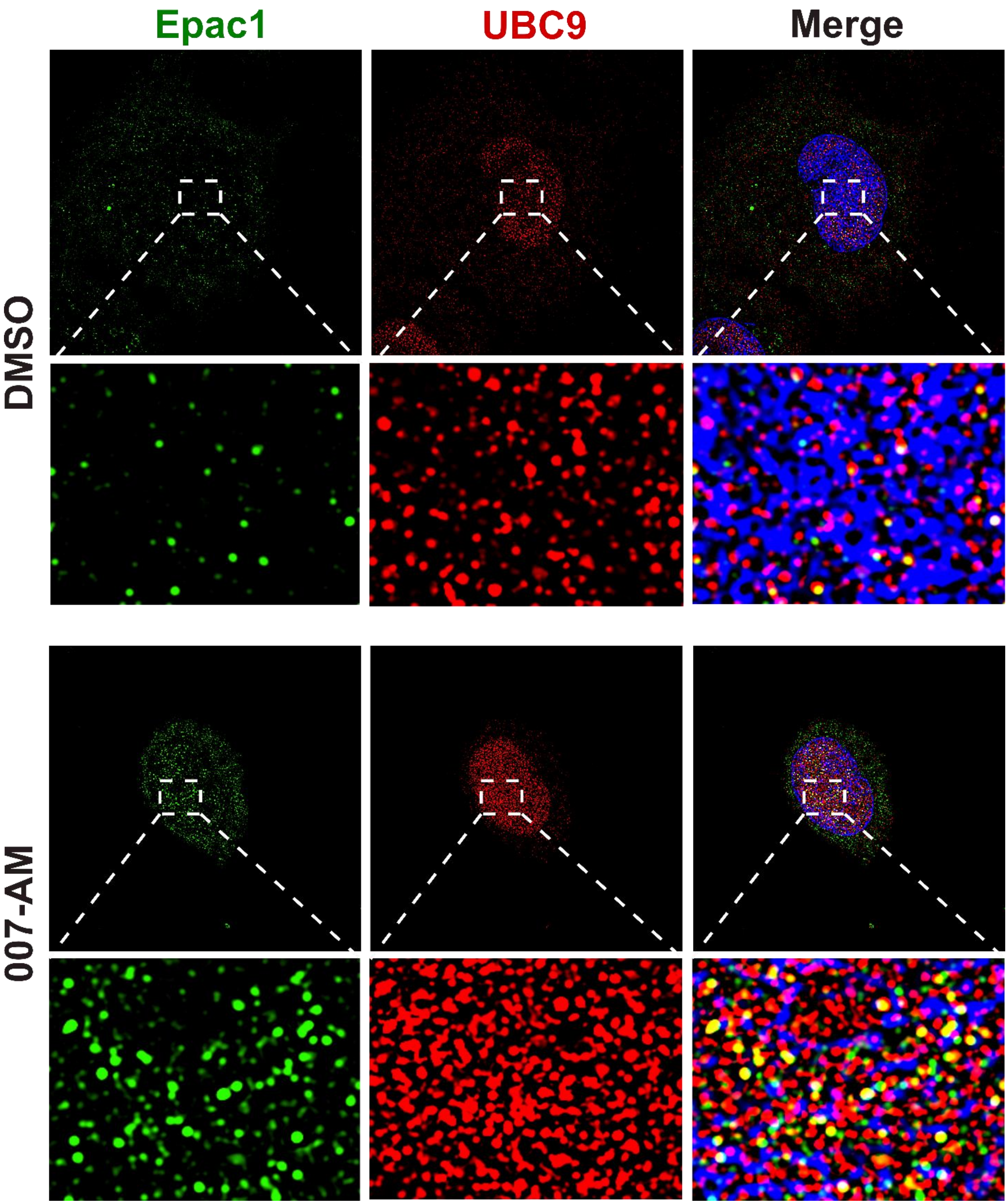


**Figure S5. Epac1 activation promotes the formation of Epac1 and UBC9 nuclear condensates and their colocalization.** SIM immunofluorescence images of endogenous Epac1 (green) and UBC9 (red) probed by anti-Epac1 (SC-28366), and UBC9 (Cell signaling, #4786) antibodies in control (DMSO) and 007-AM treated HUVEC cells.

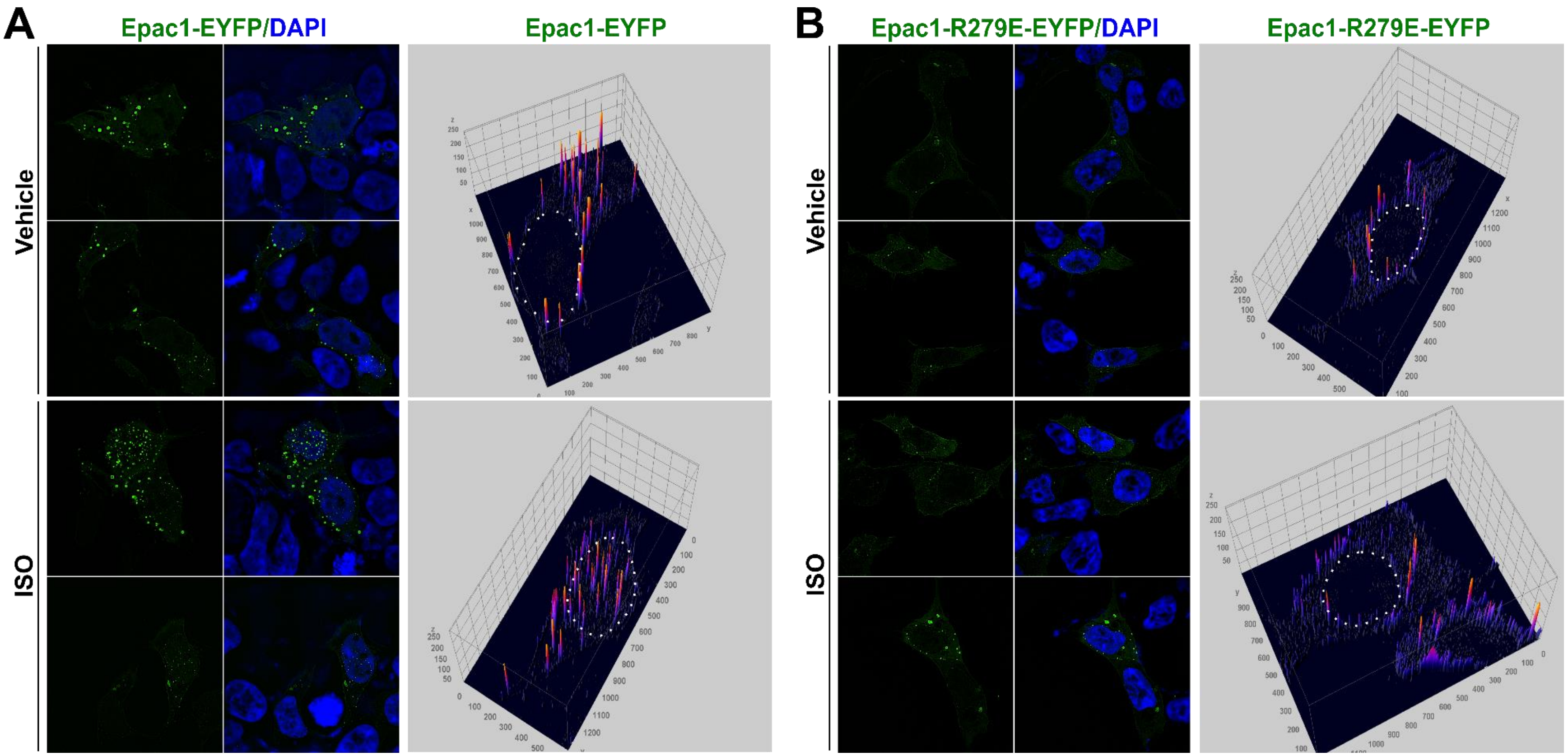


**Figure S6. ISO promotes the formation of cAMP-dependent Epac1 nuclear condensates.** Confocal fluorescence images of HEK293 cells expressing Epac1-EYFP (A) or Epac1-R279E-EYFP (B) in response to ISO (10 µM, 7 min) treatment. Dotted circles highlight the cell nuclei in the 3D contour plots.

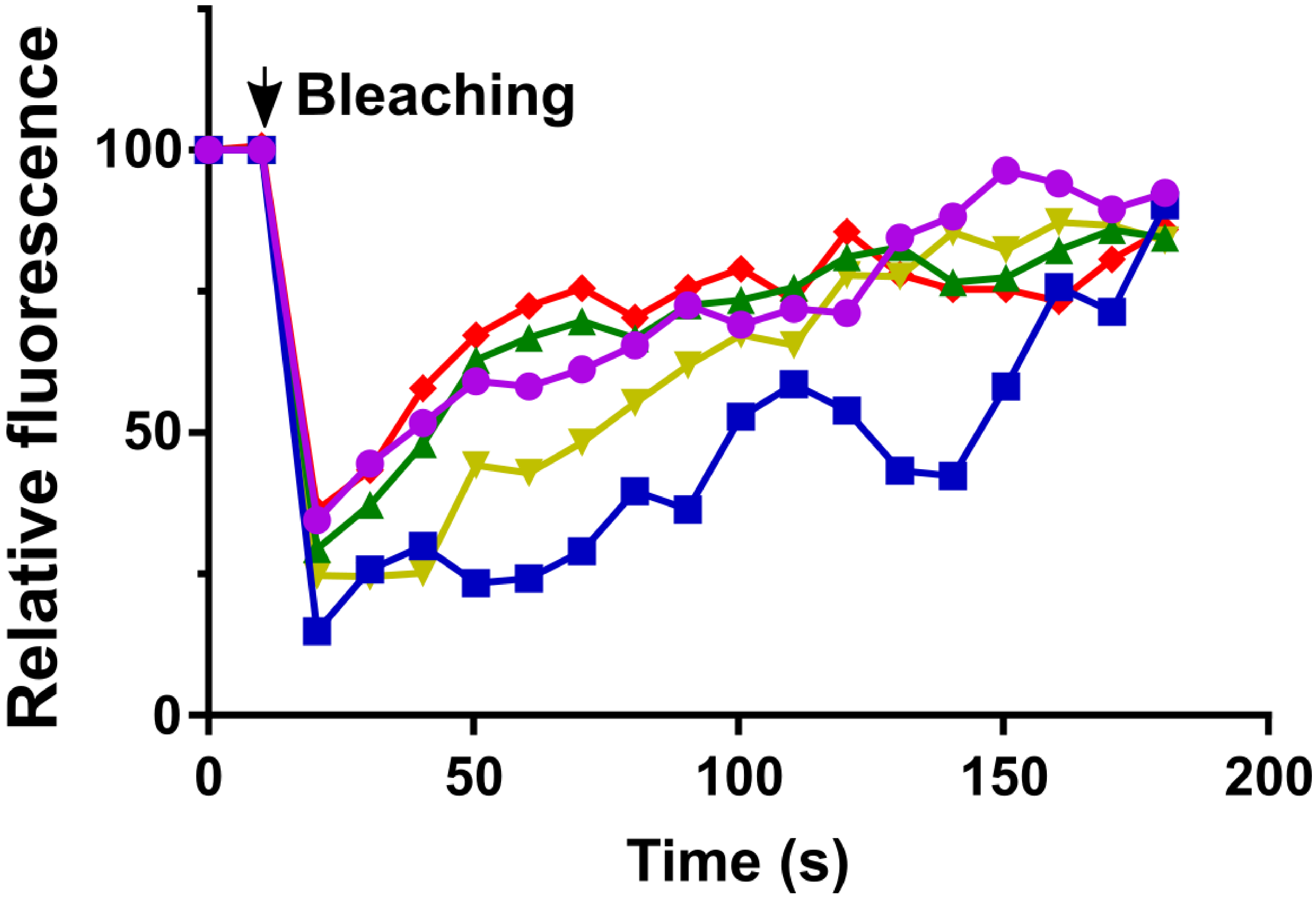


**Figure S7. FRAP analysis of Epac1-EYFP nuclear condensates.** Time-lapse traces of relative maximal fluorescence intensity of individual Epac1-EYFP nuclear condensates before and after photobleaching. Each curve represents data from one Epac1-EYFP nuclear condensate.

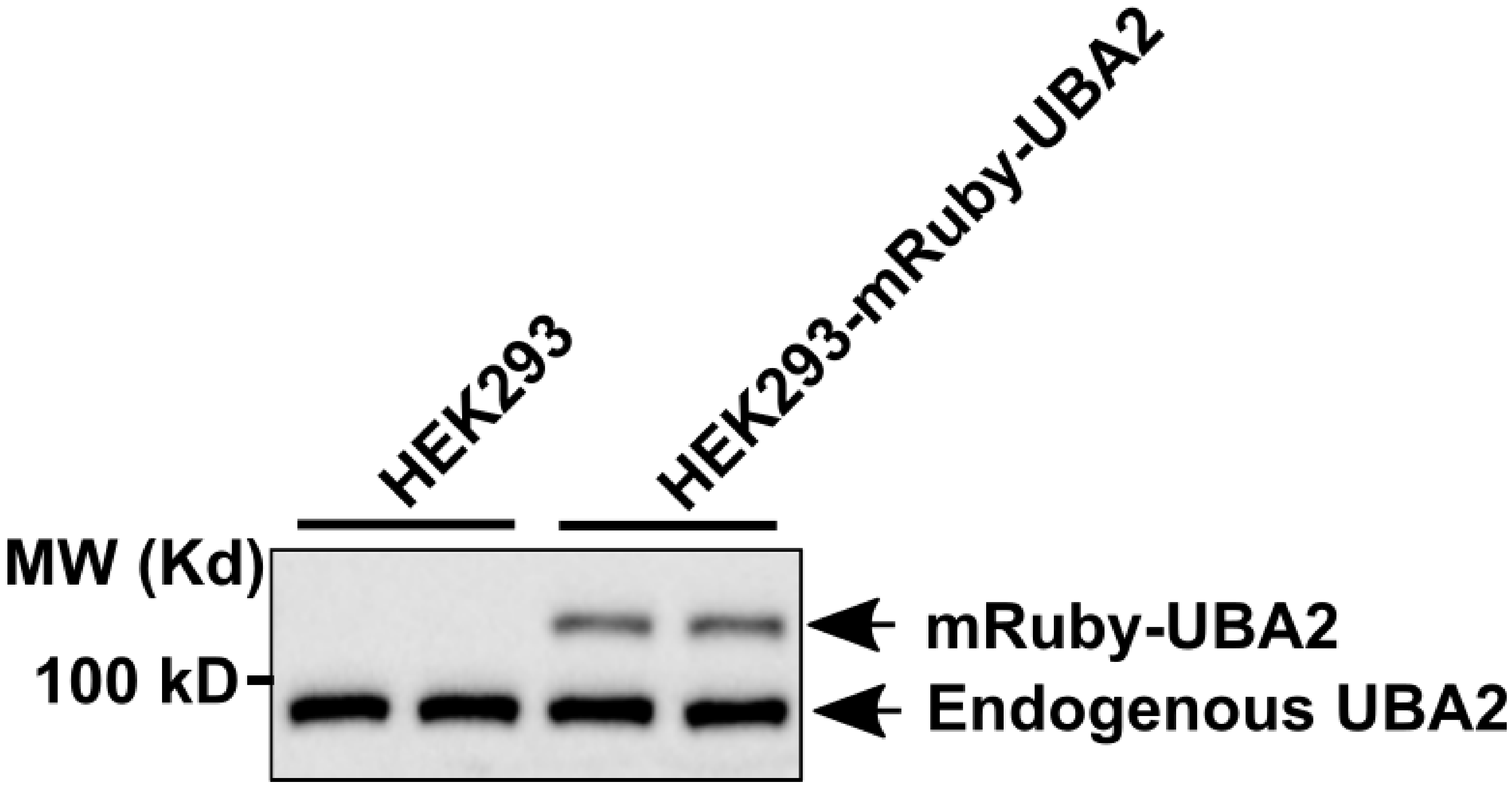


**Figure S8. Expression levels of endogenous UBA2 and ectopic mRuby-UBA2 in HEK293 cells.** Cellular UBA2 and mRuby-UBA2 levels were probed by immunoblotting using anti-UBA2 antibody.

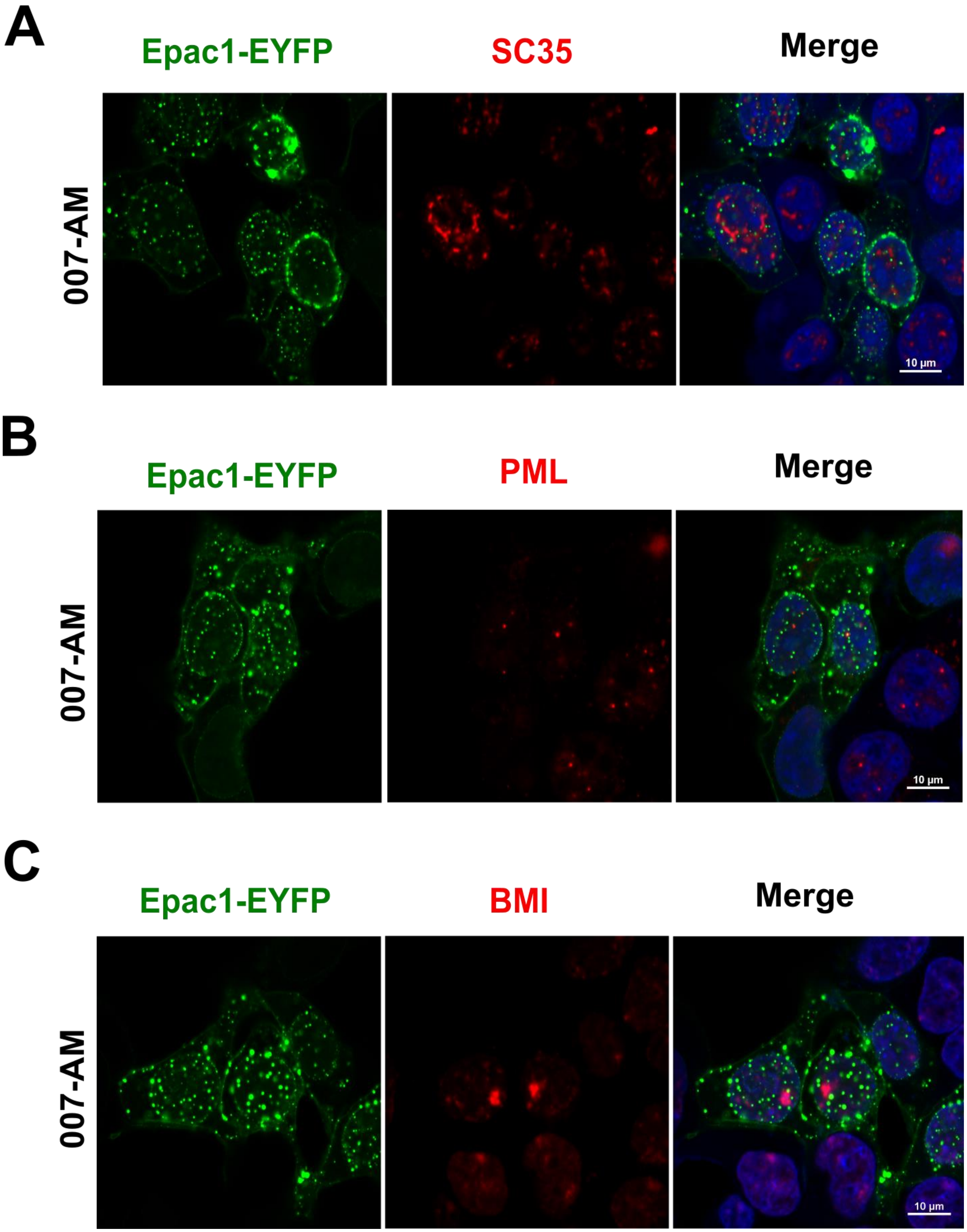


**Figure S9. Co-localization analyses of Epac1-EYFP nuclear condensates and common nuclear condensate markers.** Confocal fluorescence images of HEK293 cells expressing Epac1-EYFP co-stained with SG35 (A), PML (B) or BMI (C) in response to 007-AM.

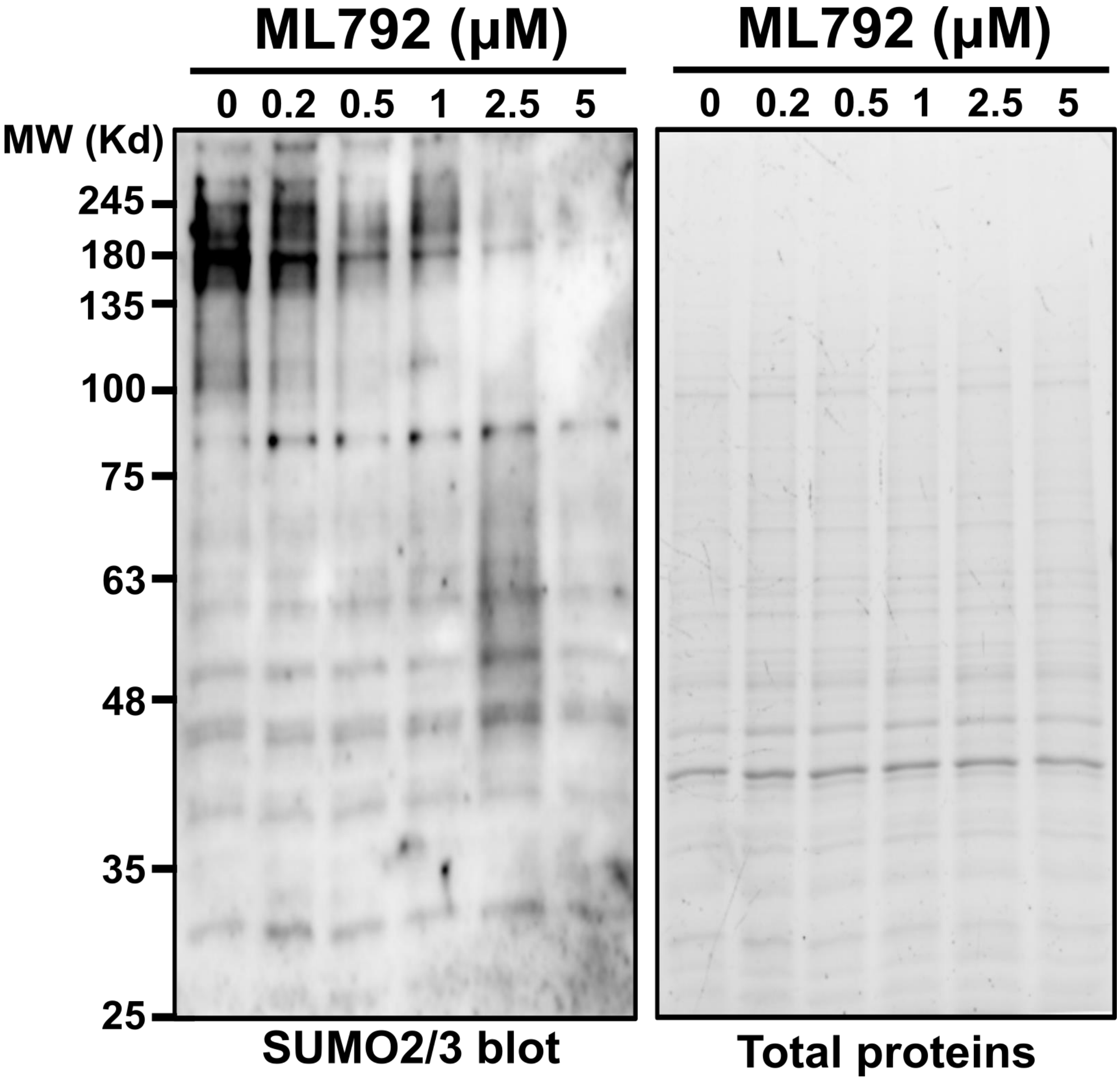


**Figure S10. UBA2 specific inhibitor ML792 blocks cellular SUMOylation in BMDMs.** Cellular SUMOylation was probed by anti-SUMO2/3 antibody in WT BMDMs treated with various concentration of ML792.
